# Sensory neuron lineage mapping and manipulation in the *Drosophila olfactory* system

**DOI:** 10.1101/312074

**Authors:** Phing Chian Chai, Steeve Cruchet, Leonore Wigger, Richard Benton

## Abstract

Nervous systems exhibit myriad cell types, but understanding how this diversity arises is hampered by the difficulty to visualize and genetically-interrogate specific lineages, especially at early developmental stages prior to expression of unique molecular markers. Here, we use a genetic immortalization method to analyze the development of sensory neuron lineages in the *Drosophila* olfactory system, from their origin to terminal differentiation. We apply this approach to first define a fate map of all olfactory lineages and refine the model of temporal patterns of lineage divisions. Taking advantage of a selective marker for the lineage that gives rise to Or67d pheromone-sensing neurons and a genome-wide transcription factor RNAi screen, we identify the spatial and temporal requirements for Pointed, an ETS family member, in this developmental pathway. Transcriptomic analysis of wild-type and Pointed-depleted olfactory tissue reveals a universal requirement for this factor as a switch-like determinant of fates in these sensory lineages.

## Introduction

Nervous systems are composed of an enormous number of cell types of diverse structural and functional properties. While the cataloguing of cell populations is advancing rapidly through single-cell sequencing approaches (Cuevas-Diaz Duran et al., 2017; Poulin et al., 2016), the genesis of most cells is poorly understood. This ignorance limits our appreciation of the relationships between the developmental trajectories, mature connectivity and functions of different cell types, as well as how genetic and environmental factors regulate the plasticity of neurodevelopment both within the lifetime of an individual and during evolution.

Tracing the development of neurons *in situ* from birth to terminal differentiation is a formidable challenge, as this process can occur over a long time period (e.g., days to weeks), and across disparate sites within the developing animal. Direct visual analyses are only practical for numerically simple (and largely transparent) nervous systems, such as *C. elegans* (Sulston et al., 1983). In more complex animals, labeling of neural lineages with dyes or retroviruses has been valuable (Clarke, 2008; Li et al., 2018; Ma et al., 2017), though these can lack spatiotemporal resolution and sensitivity. More recently, diverse genetic tools for lineage tracing have been enormously powerful in vertebrates and invertebrates (Lee, 2014; Li et al., 2018; Ma et al., 2017). However, an important constraint of genetic-based lineage tracing is that unambiguous molecular markers for specific populations of neurons either do not exist (as neural development typically requires combinatorial codes of fate determination and guidance factors (Allan and Thor, 2015; Li et al., 2017; Seiradake et al., 2016)) or are expressed only during very narrow time windows or in late stages of differentiation (e.g., sensory receptor genes). These limitations prevent reproducible genetic access to a given lineage, which necessitates laborious random genetic labeling and screening.

In this study, we address the problem of lineage tracing and manipulation in the peripheral olfactory system of *Drosophila melanogaster.* Adult olfactory organs originate from the larval antennal imaginal discs. Within this tissue, sensory organ precursor (SOP) cells are specified, and each gives rise to a short lineage producing up to four olfactory sensory neurons (OSNs) and four support cells, which together form an individual sensillum on the antenna (Barish and Volkan, 2015; Jefferis and Hummel, 2006; Vosshall and Stocker, 2007). Approximately 50 classes of OSNs exist, defined by their expression of a specific Odorant Receptor (Or) or Ionotropic Receptor (Ir) (Joseph and Carlson, 2015; Rytz et al., 2013). These OSNs are organized in stereotyped combinations within >400 antennal sensilla, which belong to several morphological subclasses: antennal basiconic/trichoid/intermediate (ab, at, ai) sensilla (which house OR-expressing OSNs) and antennal coeloconic (ac) sensilla (housing mainly IR-expressing OSNs) (Couto et al., 2005; Silbering et al., 2011; Vosshall and Stocker, 2007). The axons of OSNs expressing the same receptor converge onto a discrete glomerulus within the primary olfactory center (antennal lobe) in the brain (Couto et al., 2005; Fishilevich and Vosshall, 2005; Grabe et al., 2014; Silbering et al., 2011). Here, they synapse with projection neurons (PNs) – which arise independently from long lineages of central brain neuroblasts (Lee, 2017; Yu et al., 2010) – that carry olfactory signals to higher brain centers (Grabe and Sachse, 2018; Jefferis and Hummel, 2006).

Despite the extensive knowledge of the anatomy and function of the olfactory system (Grabe and Sachse, 2018; Joseph and Carlson, 2015; Mansourian and Stensmyr, 2015), the developmental origins of distinct olfactory SOPs and how these are related to the organization of the olfactory pathways are incompletely understood (Barish and Volkan, 2015; Jefferis and Hummel, 2006). This is largely because of the lack of genetic markers that permit the tracing of a specific lineage from the birth of an SOP to the integration of the OSN into the mature olfactory circuitry. Here we develop an immortalization labeling system for OSN lineages, which uses the principles of the CONVERT technique (Yagi et al., 2010) and takes advantage of the large resources of *Drosophila* enhancer-GAL4 driver lines for selective and reproducible genetic marking of cell subpopulations (Jenett et al., 2012; Jory et al., 2012). This approach permits us to, first, chart an olfactory fate map in the antennal disc, second, visualize, for the first time, an entire olfactory sensory lineage and, third, through a genome-wide RNAi screen of transcription factors, characterize the role of a novel molecular determinant of OSN lineage development.

## Results

### A genetic immortalization labeling system for olfactory sensory neuron lineages

We immortalized the expression properties of antennal disc-expressed GAL4 drivers within a narrow time window spanning SOP specification through three sequential events (Figure 1A): (i) temporally-controlled heat-inactivation of GAL80^ts^, a thermosensitive inhibitor of GAL4, (ii) GAL4 induction of Flippase-mediated recombination and activation of a LexA driver, (iii) stable LexA-dependent expression of a GFP reporter in the labeled SOPs and, critically, all their descendants.

**Figure 1.**
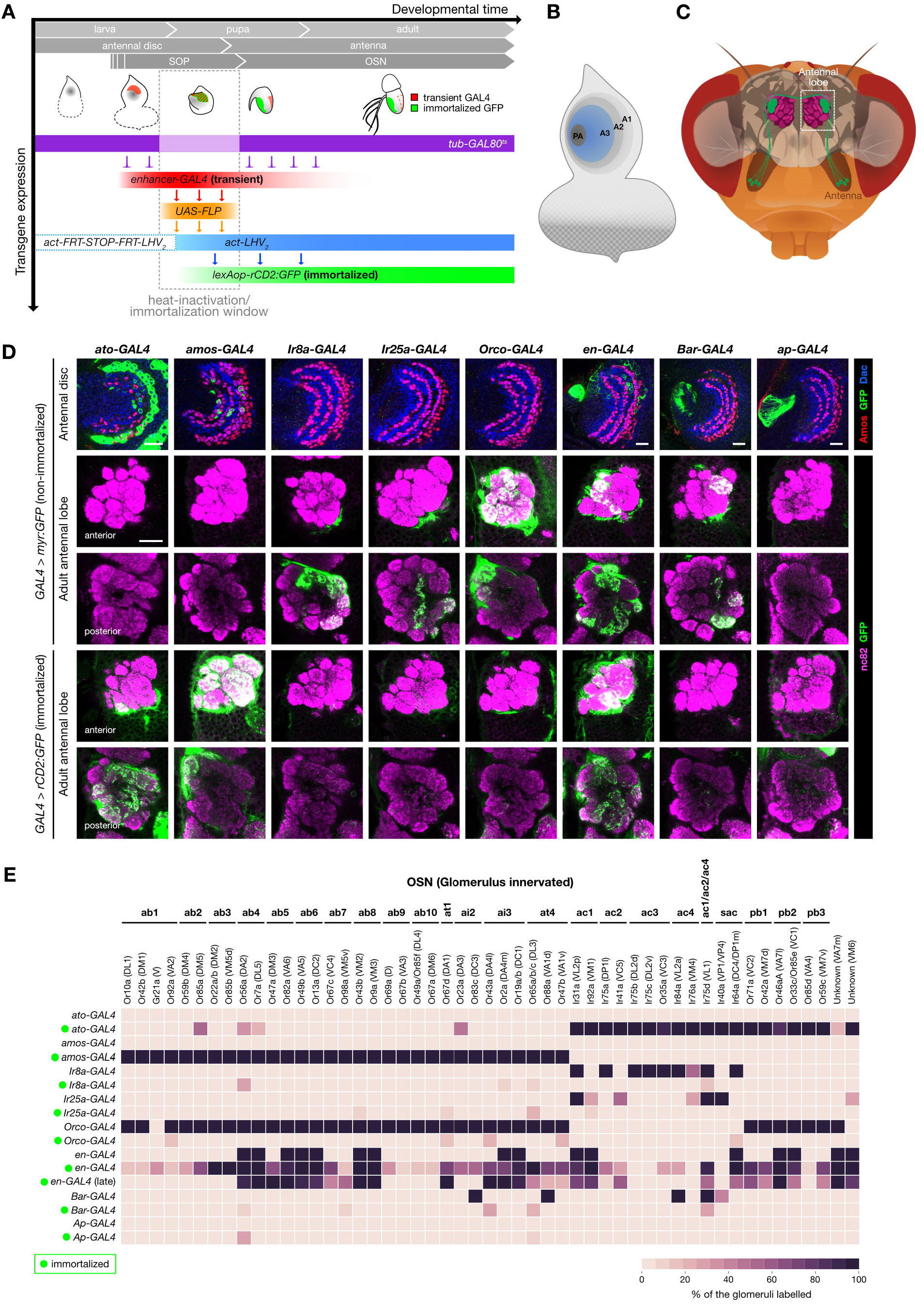
A genetic immortalization labeling system for OSN lineages. (A) Time course of the development of the peripheral olfactory system illustrating the principle of the genetic immortalization strategy. The transient expression of an *enhancer-GAL4* (red) is immortalized in a subset of SOPs during their specification at the early pupal stage, via sequential temporally-defined GAL80^ts^ heat-inactivation, Flippase (FLP)-mediated recombination, and *act-LHV2* cassette activation, to induce the permanent expression of membrane-bound myr:GFP in the SOPs and their progeny OSNs. Other cells that express the *enhancer-GAL4* outside the immortalization time window will not be labeled. (B) Schematic of the larval eye-antennal imaginal disc, illustrating the A3 region (blue), where olfactory SOPs develop. The presumptive arista (PA) – a feather-like, non-olfactory sensory structure on the antenna – forms from the center of the disc. (C) Schematic of the *Drosophila* head, illustrating a single population of OSNs expressing the same olfactory receptor (green), which project their axons from the antenna at the periphery towards a unique glomerulus of the antennal lobe in the brain (dashed box). (D) Row 1: non-immortalized *enhancer-GAL4* driven expression of myr:GFP (α-GFP; green) in 2 h APF antennal discs (except for ato-GAL4 and *amos-* GAL4, which show 4 h APF discs, due to the delayed expression of these drivers). The discs are colabeled with α-Amos (red) and α-Dac (blue) to provide spatial landmarks. Rows 2-3: non-immortalized *enhancer-GAL4* driven expression of myr:GFP (α-GFP; green) in the axon termini of OSNs innervating the right antennal lobe of the adult. The glomerular structure of the lobe is visualized with α-Bruchpilot (nc82, magenta); anterior and posterior focal planes (representing single optical slices) are presented to show the majority of glomeruli. Row 4-5: immortalized *enhancer-GAL4* driven expression of rCD2:GFP (α-GFP; green) in OSN axons, presented as in Rows 2-3. The immortalization time window was 4 h BPF-20 h APF, during which the SOPs undergo maturation (Figure 1A). Note that the immortalized enhancer-GAL4s (rows 4-5) label different subsets of OSNs from their non-immortalized counterparts (rows 2-3). Scale bars = 20 μm. The genotypes of the flies used in the figures are listed in Table S1. (E) Heatmap showing the frequency that a given antennal lobe glomerulus is innervated by OSN axons labeled by a GFP reporter (either myr:GFP or rCD2:GFP) under the control of non-immortalized or immortalized (green dot) *enhancer*-GAL4s. Glomeruli are organized by the compartmentalization of the corresponding OSNs in different sensilla classes (sac = sacculus sensilla; pb = maxillary palp basiconic sensilla; see text for other abbreviations). 7-10 antennal lobes were examined per line per condition. Immortalization was performed during the same time window (4 h BPF-20 h APF) for all drivers except for “late” en-GAL4 immortalization (9 h APF-39 h APF).

To ensure the efficacy of this system, we first compared GFP signals from “non-immortalized” and “immortalized” GAL4 drivers for two antennal proneural genes: *atonal* (ato) and *absent MD neurons and olfactory sensilla (amos),* which are expressed in SOPs that form ac and ab/at/ai sensilla, respectively (Goulding et al., 2000; Gupta and Rodrigues, 1997). In antennal discs, we focused on the proximal region (A3), labeled by patterning determinant Dachshund (Dong et al., 2002) where the olfactory SOPs are located (Figure 1B). In adults, we identified GFP-labeled OSNs based upon their glomerular innervations in the brain (Figure 1C).

At 4 h after puparium formation (APF), when SOP specification is occurring (Figure 1A), *ato-GAL4* labels many SOPs (Figure 1D), but as expression of *ato* is down-regulated by 12 h APF (prior to SOP division and neuron differentiation) (Gupta and Rodrigues, 1997), the non-immortalized driver does not label any OSNs (Figure 1D-E). By contrast, immortalized *ato-GAL4* labels OSNs in all ato-dependent sensilla (i.e., ac1-4, as well as those in the antennal sacculus and an independent olfactory organ, the maxillary palp). Similarly, non-immortalized *amos-GAL4* was detected in the disc but not mature OSNs, but when this driver was immortalized, GFP was detected in all OSNs from ab, at and ai sensilla (Figure 1D-E).

We next tested drivers for three olfactory receptor co-receptor genes *(Ir8a-GAL4, Ir25a-GAL4* and *Orco-GAL4),* which are not expressed in the disc, and only become active during later stages of neuron differentiation in many different populations of OSNs (Figure 1D-E) (Abuin et al., 2011; Larsson et al., 2004). If these drivers were immortalized during early SOP development (when they are inactive), no OSN GFP labeling was observed (Figure 1D-E).

Finally, we asked if the immortalization system could capture GAL4 driver expression that is unstable and/or transient. We chose *engrailed (en)-GAL4* because the cellular expression of *en* is highly dynamic at early pupal stages (up to 9 h APF) before stabilizing in progenitor cells (Song et al., 2012). *en-GAL4* is expressed in a large zone of the antennal disc at 2 h APF, but is restricted to just 16 OSN classes in the adult (Figure 1D-E). We immortalized this driver in either “early” (4 h before puparium formation (BPF)-20 h APF) or “late” (9-39 h APF) time windows. Early immortalization led to GFP labeling of most OSN classes, consistent with the extensive *en* expression in SOPs in early pupae (Figure 1D-E). Late immortalization led to restriction of labeling to fewer glomeruli, approaching the number labeled by the non-immortalized driver, suggesting this time window reflects *en* expression once it has largely stabilized into the final pattern observed in adults (Figure 1D-E).

Together, these results indicate that the immortalization strategy effectively captures and preserves GAL4 driver expression specifically during a desired developmental time window to relate early expression patterns in disc SOPs to the OSN lineages that arise from these precursors.

### A fate map of olfactory sensory organs precursors

A previous analysis (Li et al., 2016) proposed a fate map of SOPs in the antennal disc (i.e., defining which OSN classes precursors in different regions give rise to). This map was created, in part, by relating the endogenous spatial expression of various patterning factors in the disc to the OSN classes labeled by transgenic drivers for these same factors in later stages. Such an approach assumes that these patterning factors do not change their expression during SOP development. This assumption, however, is not necessarily correct, as indicated by the observation that drivers for patterning factors often label only a subset of OSNs within the same sensillum (i.e., which should derive from the *same* SOP) (Li et al., 2016).

To explore this issue further, we re-examined the expression of two of the previously-used drivers, *Bar-GAL4* and *apterous (ap)-GAL4.* In the antennal disc, both of these drivers are expressed in the presumptive arista (PA) zone (Figure 1B and Figure 1D). Previous analyses showed that in adults *Bar-GAL4* labels three OSN populations (VA1d/Or88a, VL2a/Ir84a and VL1/Ir75d) while *ap-GAL4* labels six populations (DA3/Or23a, VA1d/Or88a, DL3/Or65a/b/c, DM4/Or59b, DL5/Or7a, VM2/Or43b, and VL2p/Ir31a) in pupae (but not adults) (Li et al., 2016). These observations, together with loss- and gain-of-function mutational analyses, led to the proposition that SOPs for the corresponding sensilla (ab2, ab4, ab8, at2, at4, ac1, ac2, and ac4) all lie within the PA (Li et al., 2016). By contrast, we found that neither immortalized *Bar-GAL4* nor immortalized *ap-GAL4* consistently label any antennal lobe glomeruli (Figure 1D-E), and the only GFP-positive neurons we detect are located in the arista (Figure S1A-B). These observations argue that the OSN expression of the non-immortalized *GAL4* drivers reflects late expression patterns unrelated to their expression in the disc, and that the PA contains SOPs only for the arista, as previously proposed (Haynie and Bryant, 1986), but not any olfactory sensilla.

To establish an olfactory SOP fate map with our immortalization approach, we first partitioned the A3 domain of the antennal disc into eight concentric arcs (1-8), using Amos and Dac to delineate their boundaries (Figure 2A). Dac is expressed broadly in A3, forming a broad, curved zone with strong central expression and weaker expression at both dorsal and ventral edges. Amos is expressed in SOPs that form three parallel narrow bands (zur Lage et al., 2003). At 2 h APF, the SOPs within the two inner bands (arcs 2 and 4) co-express high levels of Dac; the wider outermost band overlapped with the strong-to-weak transition border of Dac expression, allowing us to subdivide it into two adjacent arcs (6 and 7). The Dac-positive, Amos-negative regions constitute the remaining four arcs (1, 3, 5 and 8) (Figure 2A). These regions are presumably occupied by Ato-positive SOPs because they flank and intercalate with Amos-expressing SOPs (Figure 2B). In addition, to increase the resolution of the spatial map, we divided each arc along the depth of antennal disc arbitrarily into superficial and deep layers (Figure 2C).

**Figure 2.**
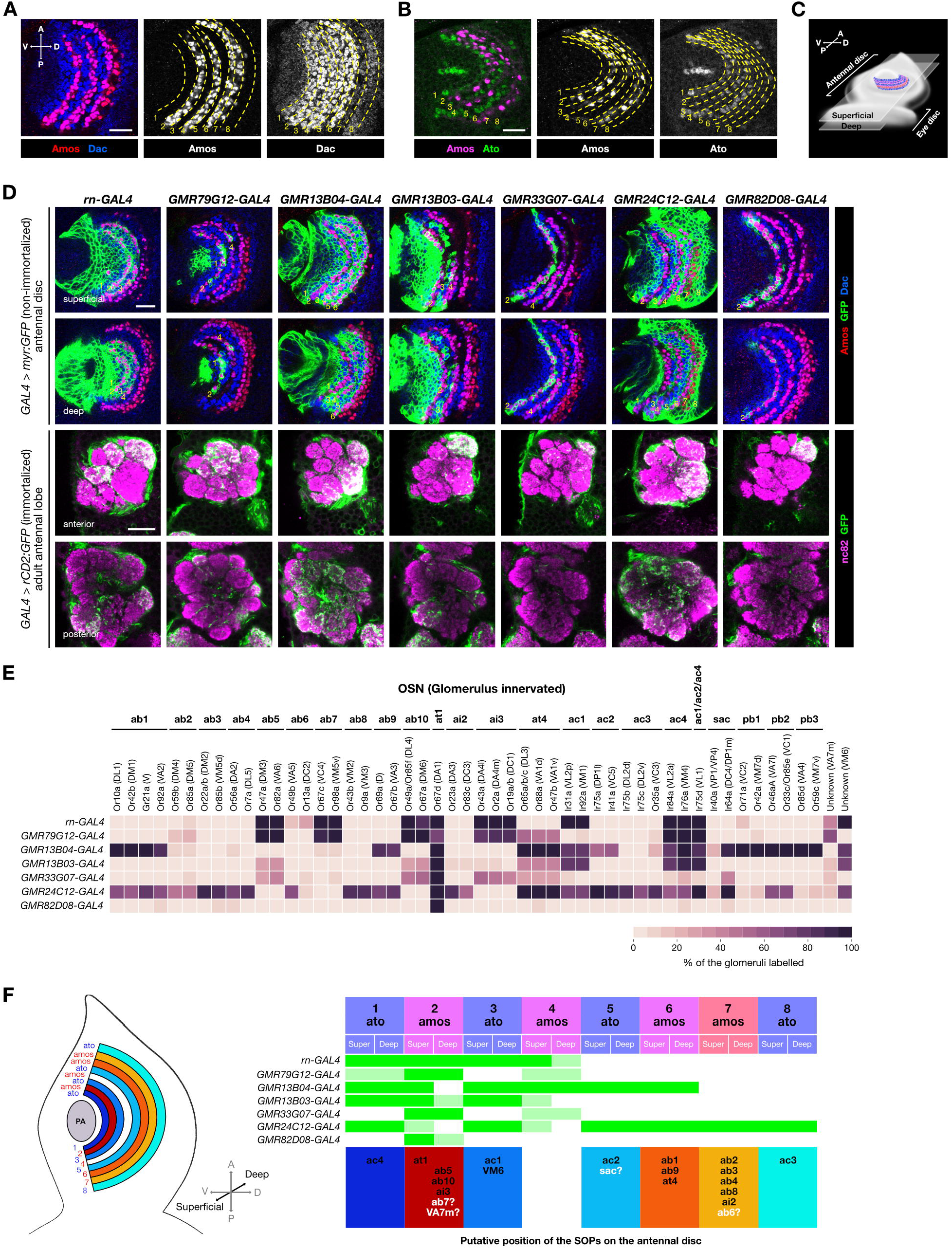
An olfactory SOP fate map. (A) Immunolabeling of a 2 h APF antennal disc (A3 region; Figure 1B) with α-Amos (red) and α-Dac (blue), whose expression boundaries allow the definition of eight distinct concentric arcs. Scale bar = 20 μm in this and other panels. (B) Immunolabeling of 4 h APF antennal disc with α-Amos (magenta) and α-Ato (green) showing that arcs 1, 3, 5 and 8 are occupied by SOPs for ac sensilla (Ato-positive) while arcs 2, 4, 6 and 7 are occupied by SOPs for ab, ai and at sensilla (Amos-positive). (C) Schematic showing the location of the arcs (indicated by the SOPs with the same color profile as in (A)) on the antennal disc to illustrate the position of the superficial and deep layers. (D) Rows 1-2: non-immortalized *enhancer-GAL4* driven expression of myr:GFP (α-GFP; green) in 2 h APF antennal discs. The discs are colabeled with-Amos (red) and -Dac (blue) to reveal GFP expression relative to the concentric arcs (yellow numbers). Images of superficial and deep layers of the disc are shown. Rows 3-4: immortalized *enhancer-* GAL4 driven expression of rCD2:GFP (α-GFP; green) in the axon termini of OSNs innervating the antennal lobe of the adult (presented as in Figure 1D). The immortalization time window was 4 h BPF-20 h APF. (E) Heatmap showing the frequency that a given antennal lobe glomerulus is innervated by OSN axons labeled by rCD2:GFP under the control of the immortalized *enhancer-GAL4* drivers shown in (D). (F) Left: schematic of the antennal disc showing the Ato and Amos-positive arcs. Right: an olfactory fate map, summarizing the expression pattern in different arcs of the *enhancer*-GAL4 lines (green: strong expression/complete coverage; light green: weak expression/incomplete coverage). The birthplace of SOPs of different OSN/sensilla classes (bottom row) could be deduced from the similarity of the expression pattern of these lines in the antennal disc and the similarity of their OSN labeling patterns when immortalized during SOP development. Black labels: high certainty of SOP assignment; white letters with question mark: ambiguous assignment.

To genetically label different subsets of SOPs, we screened images of >500 antennal disc GAL4 drivers (Jory et al., 2012), and identified 25 that exhibited restricted expression in the A3 region. Of these, 19 were subsequently eliminated as preliminary immortalization analysis gave inconsistent expression patterns suggesting these had unstable disc expression (data not shown). The expression of the remaining six GAL4 lines, along with a driver for *rotund* (*rn*) (a patterning factor with stable expression (Li et al., 2016; Li et al., 2013)), was examined within the spatially-partitioned A3 domain at 2 h APF (Figure 2D). Despite the limited number of lines, every arc on the map could be uniquely identified based on the combinatorial expression patterns of these drivers (Figure 2F). Superficial and deep layers were only resolved for arcs 2 and 4 (Figure 2F).

We next immortalized these seven GAL4 drivers and examined the identities of the labeled glomeruli (Figure 2D-E). Although these drivers also label some cells in the central brain (data not shown), antennal deafferentation experiments confirmed that the specific glomerular signals were entirely due to the contribution of OSNs (Figure S2A). While we assume that most of these OSN classes originate from SOPs within the corresponding GAL4 expression zone observed at 2 h APF, we could not definitively exclude that some OSNs might originate from other, “out-of-zone” SOPs that express GAL4 at earlier or later time-points during the immortalization window (4 h BPF-20 h APF). However, given the stochastic nature of the labeling method, we suspected such out-of-zone SOPs would lead to less frequent GFP labeling than those SOPs that we know robustly express the driver at 2 APF. By comparing the GAL4 expression patterns on the SOP spatial map and GFP-labeled OSNs after immortalization, we were able to assign the SOPs for at least 16/18 antennal olfactory sensilla classes to specific arcs (Figure 2F). For example, all OSN classes within ab1, ab9 and at4 sensilla are labeled at high frequency with the immortalized *GMR13B04-GAL4* and *GMR24C12-GAL4* lines, allowing us to infer that SOPs for these three sensilla lie within arc 6, as this is the only Amos-positive arc where only these two drivers are expressed (Figure 2E-F). The assignment of at4 to this arc remains less conclusive, because OSNs in this sensillum type are also labeled, albeit much less frequently, by other immortalized GAL4 drivers OSNs; we hypothesize that this is due to transient expression of these drivers in this arc during the immortalization window.

### Neuroanatomical correlates of the SOP fate map

We asked whether the position of SOPs on the array of antennal disc arcs has any relationship with the properties of these circuits in the adult. In contrast to the earlier fate map (Li et al., 2016), we find that individual subtypes of ac, ai and at sensilla originate from distinct arcs (Figure 2F); for the more numerous ab sensilla, several subtypes derive from the same arc (Figure 2F). Mapping of arc identity onto the spatial distribution of sensilla in the adult antenna (Grabe et al., 2016) revealed partial preservation of the concentric organization of different lineages (Figure S2B). There are, however, some exceptions (e.g., a lateral subset of ac3 sensilla), which may reflect both the distortion of the original spatial relationships of SOP lineages due to eversion of the antennal disc during metamorphosis, as well as the local dispersion of sensilla in the pupal antenna (Song et al., 2012). At the molecular level, there is no obvious relationship between receptor sequence and arc of origin of the corresponding OSN: any individual arc gives rise to OSNs expressing Ors (or Irs) that are not confined to a specific clade on phylogenetic trees of these receptor repertoires (Figure S2C).

Examination of arc identity and antennal lobe glomerular organization revealed a strong segregation of OSNs derived from Amos- and Ato-expressing arcs, as previously described (Couto et al., 2005; Silbering et al., 2011). Within these major subcategories, however, we detected no obvious clustering of glomeruli innervated by OSNs derived from the same arc (Figure S2D).

Finally, we noted a bias in SOP arc placement and the birth order of the projection neuron (PN) synaptic partners in the near-completely mapped anterodorsal PN neuroblast lineage (Yu et al., 2010): of the early (embryonic) born PNs the majority (10/13) partner with OSNs originating from more internal arcs (1-3), while of the late (larval) born PNs the majority (12/13) partner with OSNs originating from more peripheral arcs (6-8) (Figure S2E). We speculate that this pattern reflects a remnant of evolution, and propose the following model: ancestral olfactory circuits are formed from OSNs derived from internal arcs and early-born PNs. More recently-evolved olfactory pathways comprise OSNs derived from SOPs in arcs that were added externally to the array in the disc, and PNs that were generated by extension of the central neuroblast divisions.

### A genetic driver for a single OSN lineage

Amongst the lines used to generate the fate map, we further focused on *GMR82D08-GAL4.* This driver labels SOPs that exclusively produce at1 sensilla (Figure 2D-E), which contain a single Or67d-expressing OSN that projects to the DA1 glomerulus (Ha and Smith, 2006; Kurtovic et al., 2007). This reagent (hereafter, “at1 driver”) was of substantial interest in permitting developmental analysis of a specific OSN lineage. The at1 driver is relatively stably expressed in the SOPs located around arc 2 of the antennal disc from late larval until early pupal stages (Figure 3A). From 20 APF, certain cells within the newly-formed SOP-derived cell clusters start down-regulating at1 driver expression until it is no longer detectable by 48 h APF (Figure 3A). When immortalized, the at1 driver persists to label ~35 cell clusters (mean 35.4 ± 6.4 SD; n=15 antennae) in the adult antenna, of which ~33 contain a single cell that expresses *Or67d* mRNA (33.1 ± 4.1 / 35.4 ± 6.4 (93.5%) clusters) (Figure 3B). The remaining four cells in the labeled clusters correspond to sensilla support cells, as they express the odorant binding protein Lush (Xu et al., 2005) (Figure 3C-D). As there are ~60 Or67d OSNs in the adult antenna (Grabe et al., 2016), these observations indicate that the at1 driver is expressed specifically, though incompletely, in at1 SOPs.

**Figure 3.**
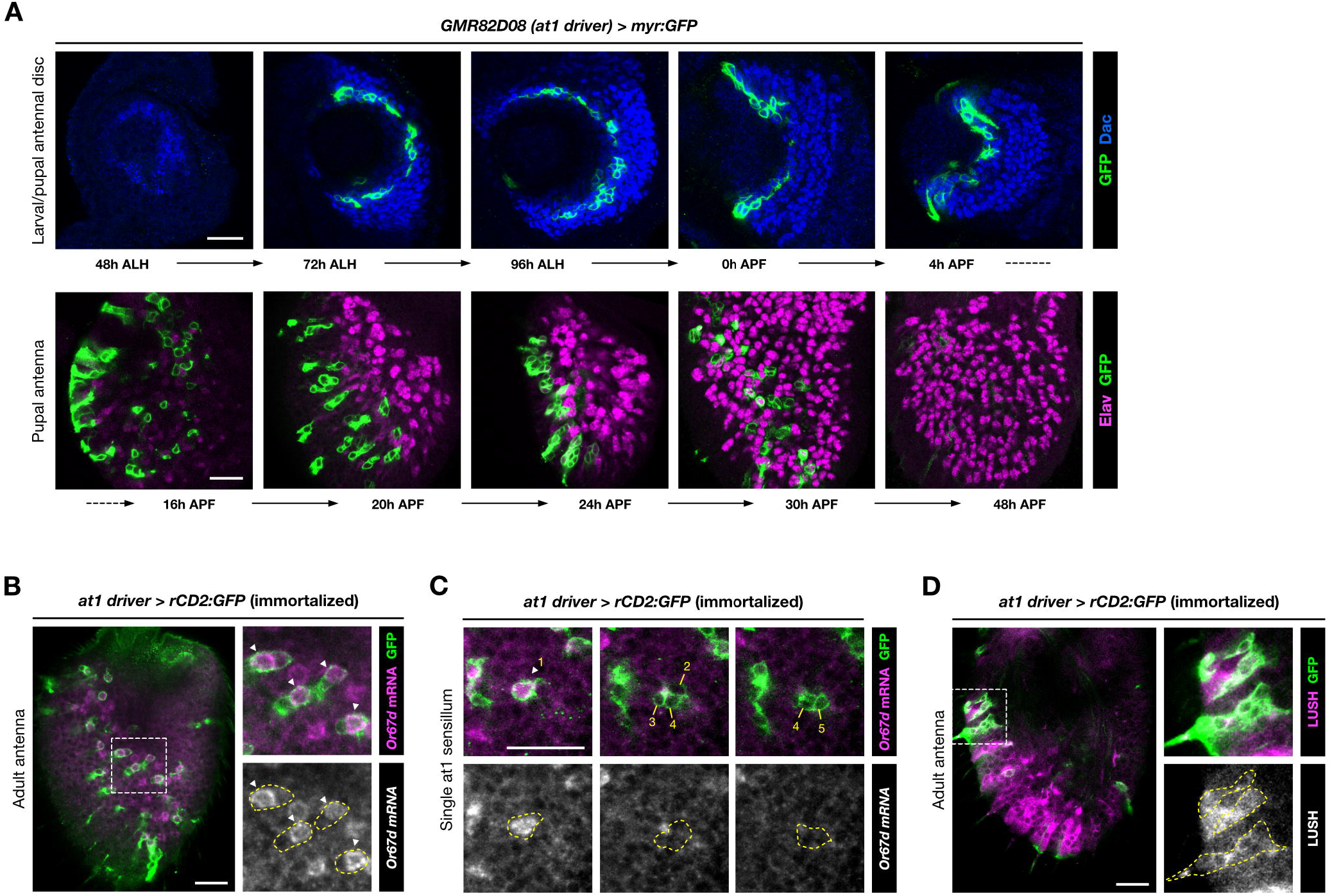
An OSN-specific lineage driver. (A) Top row: developmental expression of the non-immortalized *GMR82D08-GAL4* (hereafter, at1 driver) using a myr:GFP reporter (green) in the antennal disc SOPs (region marked by-Dac (blue)) during late larval/early pupal stages. Bottom row: the at1 driver is expressed in the daughter cells of these SOPs in the developing pupal antenna but progressively loses its expression from 20 h APF as OSNs differentiate (as visualized with the neuronal marker α-Elav (magenta)). Scale bar = 20 μm, in this and other panels. (B) Immortalization of the at1 driver reveals labeling of clusters of cells in the adult antenna by a rCD2:GFP reporter (green). Fluorescence *in situ* hybridization demonstrates that a single cell within each cluster (arrowheads in the inset images) expresses *Or67d* mRNA (magenta). (C) Representative example of a single sensillum in the adult antenna labeled by the immortalized at1 driver, viewed at three focal planes. There is a single *Or67d* mRNA-positive OSN (cell 1, arrowhead), flanked by four nonneuronal support cells (cells 2-5). (D) Sensilla cells labeled by the immortalized at1 driver lineage (α-GFP; green) also express Lush (magenta), an odorant binding protein unique trichoid sensilla support cells (Kim et al., 1998).

### Analysis of OSN lineage properties with the at1 driver

Previous analysis of olfactory lineages, using broadly-expressed enhancer trap lines for progenitors and cell division markers led to the proposal that SOPs do not divide but locally recruit other precursor cells to form a sensillum (Rodrigues and Hummel, 2008). This idea was subsequently discarded, as the use of random, clonal labeling approaches and visualization of several transcription factors allowed reconstruction of a model of SOP lineage divisions from observation of many independent clones (Endo et al., 2007; Endo et al., 2011).

An important drawback of the latter approach, however, is its lack of specificity for visualizing particular SOP subtypes. Indeed, we found that clones generated in this manner within the same time window can be both variable in cell number and expression of developmental markers (Figure S3A-B). It is impossible to distinguish whether this is because they represent distinct SOP lineages or because SOPs of the same type develop asynchronously. Using our immortalized at1 driver to visualize only this olfactory lineage, we observed that the timing of cell division and transcription factor expression in different at1 SOPs is highly synchronized (Figure 4A). This observation indicates that the random clonal approach visualizes, as expected, multiple lineages, which confounds appreciation of the precise developmental properties of a single lineage.

**Figure 4:**
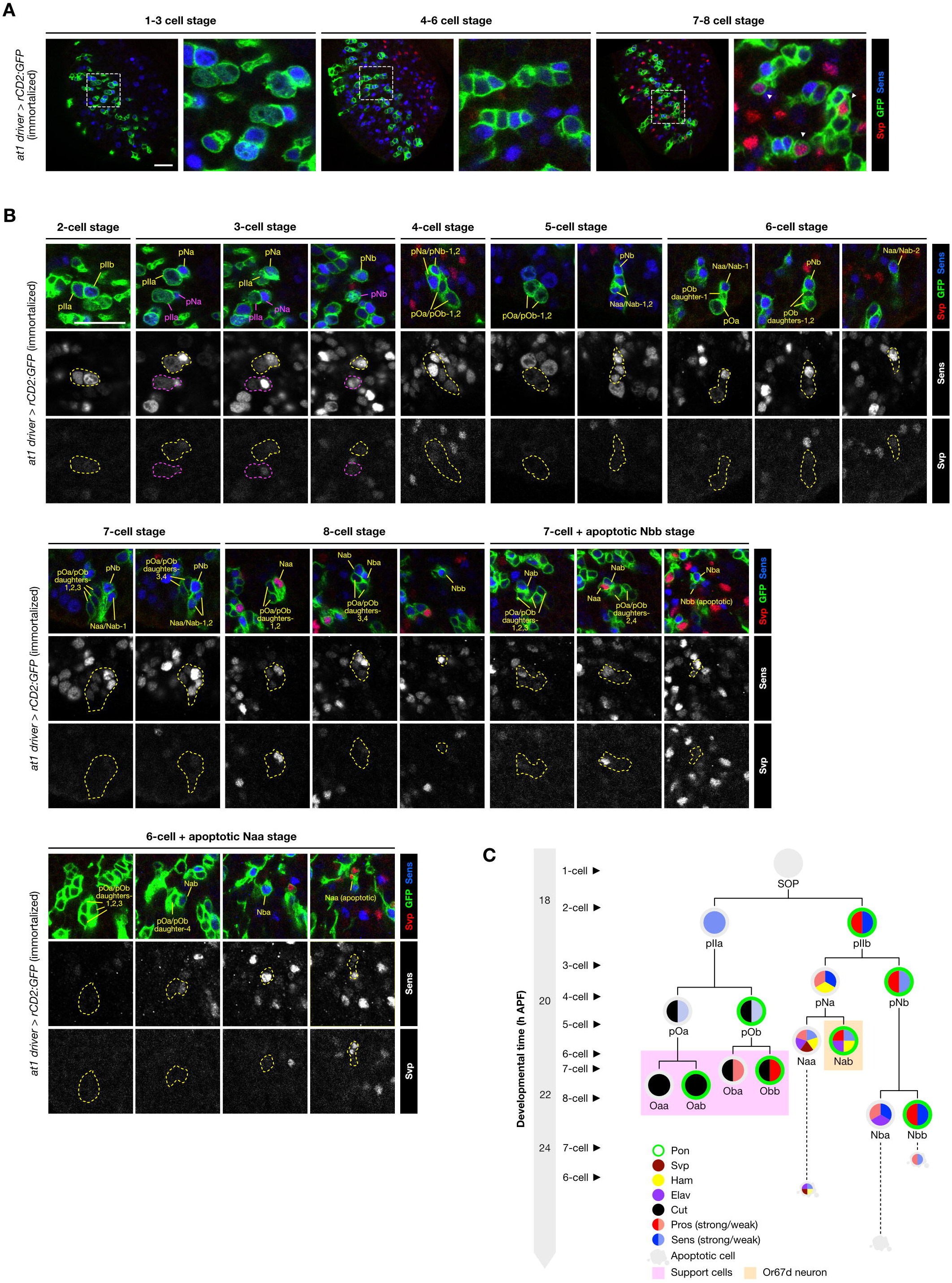
The at1 lineage is highly synchronized both in cell division sequence and molecular marker expression. (A) Full views of the immortalized at1 driver-labeled lineages (green) on the pupal antennae at three developmental time points. For each time point, all the at1 lineages have comparable numbers of daughter cells and identical Svp (red) and Sens (blue) expression patterns (shown in the magnified views of the boxed regions). Scale bar = 20 μm in this and other panels. (B) Close-up views of the SOP and its daughter cells in the at1 lineage (green) reveal a highly stereotyped series of cell divisions from 2 to 8 cells, before the Nbb and Naa cells undergo sequential apoptosis (the apoptosis of Nba cell is expected to happen after 28h APF). Most of the intermediate and terminal daughter cells in this lineage could be unambiguously identified based on the expressions of Svp (red) and Sens (blue), as well as their positions within the cluster (data not shown). Where necessary, multiple focal planes are shown to visualize all the cells in each cluster. (C) Schematic showing the division time, birth sequence and the expression pattern of common molecular markers in the at1 lineage (based also upon data from Figure S3).

We therefore reanalyzed the spatiotemporal properties of molecular marker expression throughout the at1 lineage. Observation of just two lineage markers, Senseless (Sens) and Seven-up (Svp), allowed us to establish the precise birth order of cells, which was not possible in the random labeling approach (Endo et al., 2007; Endo et al., 2011) (Figure 4B). Visualization of additional markers (Partner of Numb (Pon), Hamlet (Ham), Embryonic lethal abnormal vision (Elav), Cut (Ct), and Prospero (Pros)) permitted unambiguous determination of the identities of all intermediate and terminal daughter cells fates of this lineage (Figure 4B-C and Figure S3C-E). These comprise the Nab cell (which gives rise to the Or67d OSN (Endo et al., 2011)), the three other cells of the neuronal lineage that undergo apoptosis (Naa, Nba, Nbb), and four support cells (Oaa, Oab, Oba, Obb).

### A screen for molecules controlling at1 lineage development identifies the ETS domain transcription factor Pointed

Although several factors that function in OSN lineage-specification have been identified (reviewed in (Barish and Volkan, 2015)), our knowledge of this process – which entails the specification and coordination of olfactory receptor expression with axon guidance to a specific glomerulus – is incomplete. To identify novel molecules regulating Or67d OSN specification, we conducted a transgenic RNAi screen of 808 genes encoding candidate transcription factors, chromatin regulators and embryonic patterning genes. Knockdown of 121 genes showed various forms of antenna developmental defects (P.C.C. and R.B., unpublished data). Secondary screening with independent transgenes yielded 35 high confidence hits that gave reproducible phenotypes with at least two RNAi constructs (Figure 5A).

**Figure 5:**
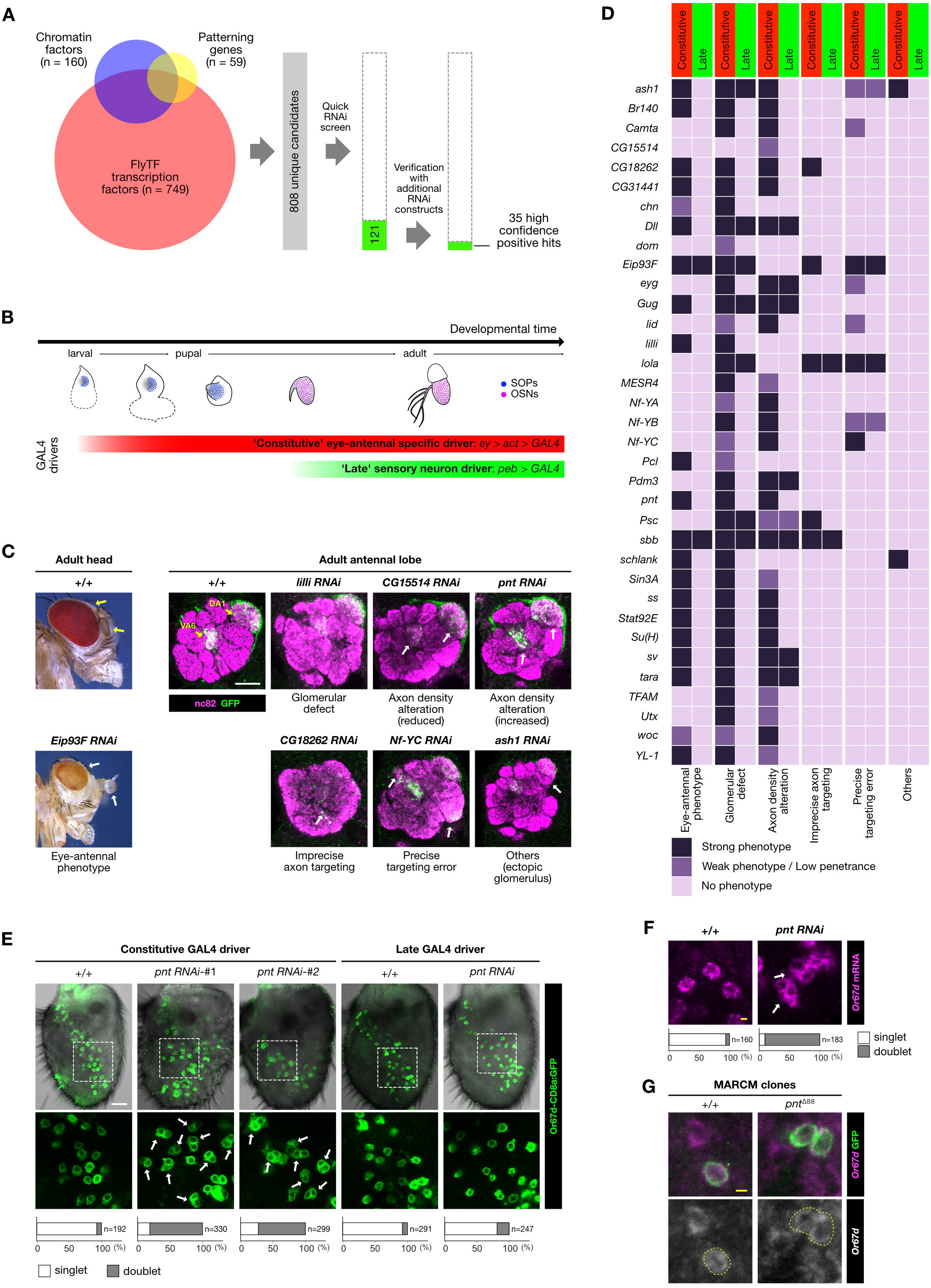
A transgenic RNAi screen for molecular determinants of at1 lineage development. (A) RNAi screen workflow: two rounds of screening, starting from 808 candidate genes [collated from (i) transcription factors (FlyTF.org (Pfreundt et al., 2010)), (ii) proteins involved in chromatin-related processes (FlyTF.org (Pfreundt et al., 2010)), and (iii) embryonic patterning genes (gap genes, pair-rule genes and segment polarity genes listed in The Interactive Fly (sdbonline.org/sites/fly/aimain/5zygotic.htm)] identified 35 genes that showed defects in antennal development upon knockdown with two or more independent transgenic RNAi constructs. (B) Temporal expression of GAL4 driver lines used for RNAi: the “constitutive” driver, comprising *ey-FLP* and an *actin-GAL4* “flip-out” cassette, is expressed specifically in the eye-antennal tissue from the second instar larval stage onwards, spanning SOP specification, divisions and OSN differentiation (red bar). The “late” *peb-GAL4* driver is expressed only after OSNs begin differentiation (green bar). (C) Phenotypic classification of the 35 high confidence screen hits when the RNAi transgenes were driven by either the constitutive or late driver. Every gene is classified under six phenotypic categories in three levels of phenotypic severity (strong, weak and none). (D) Examples of the six phenotypic categories. Left: lateral views of the fly head, with the eyes and antennae marked by arrows. When *Eip93E RNAi* is induced with the constitutive driver, there is a decoloration of the eyes, in addition to morphological defects of both eyes and antennae. Right: right antennal lobes of control and RNAi animals expressing an Or67d-CD8a:GFP reporter (α-GFP; green), which labels the axons of the Or67d and Or82a OSNs that innervate DA1 and VA6 glomeruli (yellow arrows), respectively. RNAi phenotypes of specific genes with the constitutive driver include global glomerular morphology defects, axon density alteration, imprecise axon targeting, precise targeting error and/or other defects. The phenotypic defects are indicated by the white arrows. White scale bar = 20 μm in this and other panels. (E) Phenotypes of control and *pnt* RNAi when driven with the constitutive and late drivers in antennae expressing the Or67d-CD8a:GFP reporter (green). Two independent *pnt* RNAi lines – KK100473 (#1) and JF02227 (#2), which target different genomic sequences within an exon common to all four *pnt* isoforms – result in doublets of GFP-positive neurons, compared to the predominant singlets of these neurons in controls. Late induction of RNAi produces few doublets. The lower images show the magnified views of the region within the dashed box in the upper images. Quantification of the fraction of singlets/doublets are shown in the stacked histograms at the bottom. (F) Control at1 sensilla have a single *Or67d* mRNA-expressing OSN (magenta). In *pnt* RNAi antennae, *Or67d* mRNA is detected in doublets of OSNs. Yellow scale bar = 2 μm in this and other panels. (G) Representative MARCM clones of control and *pnt* null mutant neurons, marked with CD8a:GFP (green). *Or67d* mRNA expressing OSNs (magenta) in control clones are singlets, while those in *pnt* mutant clones are doublets.

We next performed detailed analyses on these 35 genes by inducing RNAi using two GAL4 drivers with different spatiotemporal expression profiles: first, a “constitutive”, eye-antennal disc-specific driver that is active from second instar larval stage onwards (data not shown), and second, a “late” sensory neuron driver that is activated during OSN differentiation (Sweeney et al., 2007) (Figure 5B). To focus on Or67d OSN fate specification, we performed RNAi in flies expressing a Or67d:GFP reporter (this additionally labels Or82a OSNs due to ectopic transgene expression) (Couto et al., 2005). To inform both OSN fate specification and wiring, we visualized the innervation of neurons expressing this reporter in the antennal lobe, and classified the RNAi phenotypes into distinct categories (Figure 5C). While all of these genes gave strong phenotypes with the constitutive driver, only 12/35 did so with the late driver, implying that the majority is important for early developmental processes (Figure 5D). Loss of function of most of these genes led to a reduction of Or67d axon density in DA1, potentially caused by misspecification in the at1 lineage of OSN fate, receptor expression and/or OSN targeting (Figure 5C). One intriguing exception was observed in *pointed (pnt)* RNAi flies, which exhibited a higher density of Or67d OSNs innervating the antennal lobe, as revealed by an enlarged DA1 glomerulus (Figure 5C). A similar phenotype was observed for Or82a OSNs that target the VA6 glomerulus (Figure 5C).

To understand the basis of the enlarged glomerulus phenotype, we examined reporter expression in the antenna and observed a substantial increase in the number of GFP-positive OSNs (81.2±16.9 (n=12 antennae) in control animals and 177.5±33.3 (n=10 antennae) in *pnt* RNAi animals). Moreover, while the soma of the labeled OSNs in control antennae are separated from one another (“singlets”, reflecting their compartmentalization in distinct sensilla), in *pnt* RNAi antennae most of the GFP-expressing soma occur in clusters of two cells (“doublets”) (Figure 5E). This phenotype was not observed using the late driver of *pnt* RNAi (Figure 5D-E), suggesting that *pnt* regulates OSN fate prior to terminal OSN differentiation. The duplication of *Or67d* neurons in *pnt* RNAi antennae observed with the GFP reporter was validated by direct visualization of *Or67d* mRNA expression (Figure 5F). Furthermore, we confirmed this RNAi phenotype through analysis of a *pnt* loss-of-function mutation, by using MARCM (Lee and Luo, 1999) to generate GFP-labeled wild-type or *pnt* mutant OSN clones, which were examined for *Or67d* mRNA expression. All Or67d-expressing neurons in control clones (n=14) were singlets; by contrast, all *pnt* clones (n=6) contained doublets of *Or67d* mRNA-positive neurons (Figure 5G).

### Pnt is expressed dynamically within the at1 lineage

To understand the function of Pnt in the at1 lineage, we first examined the expression of this transcription factor. Because Pnt is expressed in many different cells in the pupal antenna (Figure 6A), our immortalized at1 driver line (Figure 3) was essential to follow its dynamic expression explicitly in this lineage (Figure 6A-B). Pnt is weakly, but broadly, expressed at early pupal stages (2 h APF) (Figure S4A) and becomes upregulated in the at1 SOPs just prior to their division (Figure 6A). Its expression is immediately turned off after cell division – as the daughters of this SOP (pIIa and pIIb) are devoid of Pnt immunoreactivity – and resumes in all progeny of pIIa that form support cells. In the neural sublineage derived from pIIb, Pnt is only expressed in pNa and both its daughters Naa and Nab. However, its expression is rapidly down-regulated in Nab cells, leaving Naa (marked by Svp) the only terminal cell in the neural sublineage to express this transcription factor (Figure 6A-B).

**Figure 6.**
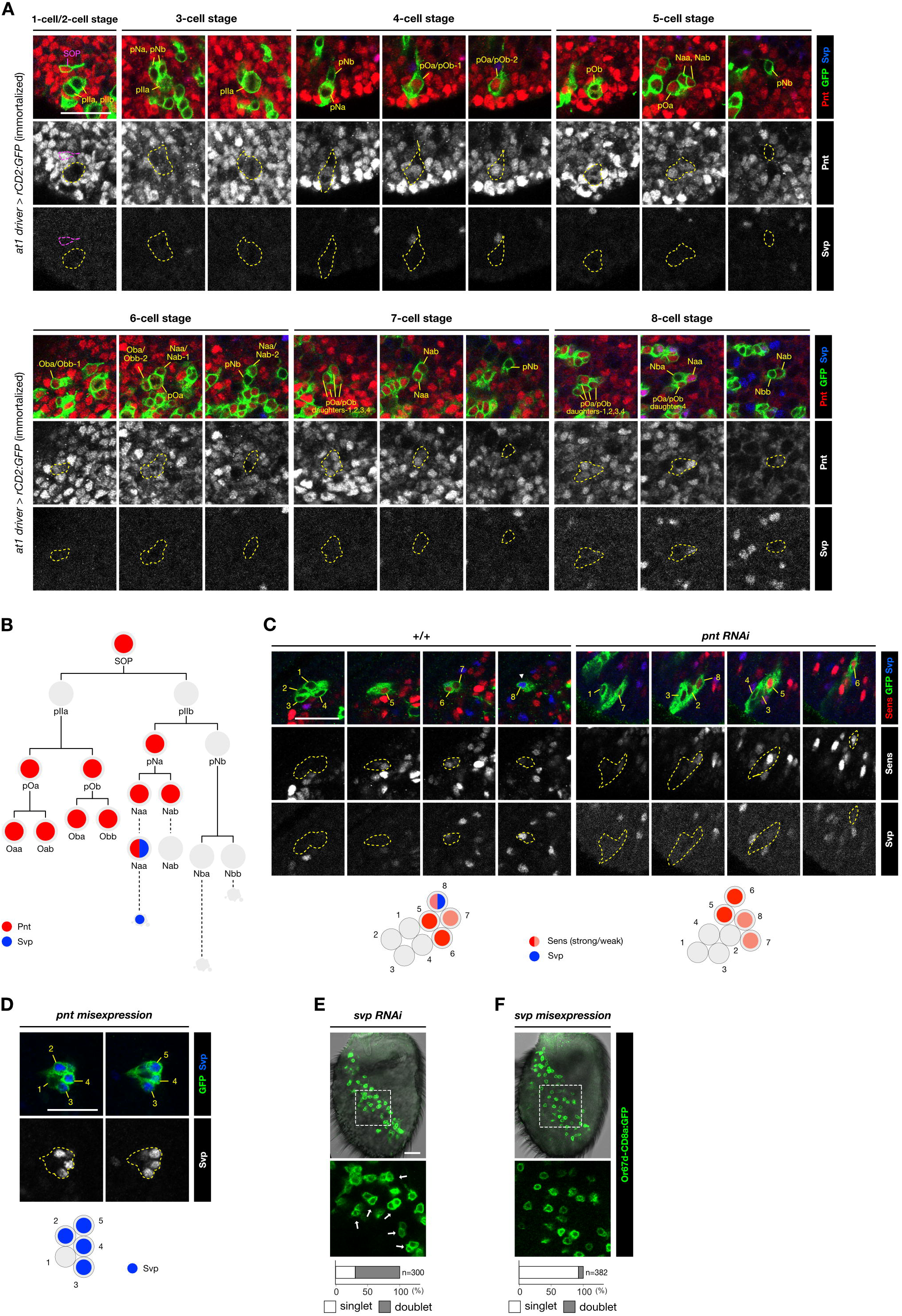
Pnt is dynamically expressed in the at1 lineage and required for Svp expression in the Naa terminal cell. (A) Expression profiles of Pnt (red) and Svp (blue) in at1 lineages visualized with the immortalized at1 driver (green). Scale bar = 20 μm in this and other panels. (B) Schematic summarizing the expression of Pnt and Svp in the at1 lineage. (C) Left: representative control 8-cell heat-shock clone (green) that shows the typical expression profile of at1 lineage: two cells with stronger Sens expression (red), and two cells with weaker Sens expression, one of which is co-expressing Svp (blue) (see Figure 4). Right: in a representative 8-cell *pnt* RNAi clone (green), Sens expression (red) is unaffected but Svp expression (blue) is absent. Schematics summarizing the expression of the markers in these clones are shown below the images. (D) Misexpression of Pnt in a 5-cell clone (green) induces ectopic Svp expression (blue) in four of the cells. (Clones do not develop beyond this stage). (E) A *svp* RNAi antenna expressing the Or67d-CD8a:GFP reporter exhibits a large fraction of doublets of GFP-positive neurons (green) (F) Misexpression of *svp* does not affect the normal specification of *Or67d-* CD8a:GFP-labeled OSNs. Two independently constructed *UAS-svp* transgenes were used in this experiment: *UAS-svp.II* (shown in this panel) and *UAS-svp1.*

### Pnt is essential for the distinction of Naa and Nab neuron fates

The complex spatiotemporal expression pattern of Pnt raised the possibility that it acts in multiple cell fate determination events in the at1 lineage to restrict the differentiation of a single Or67d OSN. To understand the genesis of the duplicated Or67d OSNs when Pnt is lost, we generated GFP-labeled wild-type or *pnt* RNAi cell clones and examined the expression of diagnostic cell markers at the 8-cell stage. Although we were unable to selectively label the at1 lineage in these experiments, we could identify putative 8-cell at1 clones by their spatial distribution in the pupal antenna (Figure 4A) and their characteristic expression of Sens, as determined by our lineage mapping (Figure 4B-C). Comparison of 8-cell wild-type and *pnt* RNAi clones revealed indistinguishable patterns of Sens and Ham expression in 4 and 2 cells, respectively (Figure 6C and Figure S4B). These observations suggest that loss of Pnt does not lead to conversion of Pnt-positive support cell lineages into neuronal fates, nor does it transform the Pnt-positive pNa lineage into the pNb lineage. By contrast, expression of Svp, which labels uniquely the Naa cell in controls, is lost in almost all (16/17) *pnt* RNAi clones examined (Figure 6C). This observation suggests a model in which Pnt functions in the Naa cell to induce Svp expression, which distinguishes its fate from the Nab cell. In the absence of Pnt, the pNa cell divides to produce two Svp-negative Nab cells, which differentiate to form the observed doublet of Or67d OSNs.

We tested this model first by asking whether Pnt was sufficient to promote Naa fate by misexpressing Pnt in SOP clones (Figure 6D). Many of the intermediate daughter cells within each SOP lineage expressed Svp, consistent with the ability of Pnt to induce this marker of Naa cells. We were unable, however, to examine later phenotypes as the ectopic expression of Pnt and/or Svp blocked lineage development at the 5-cell stage (Figure 6D). Next, we examined the role of Svp itself, which has only previously been used as a marker of Naa fate. Strikingly, *svp* RNAi in the antenna phenocopies *pnt* RNAi giving rise to doublets of Or67d OSNs (Figure 6E). However, mis-expression of *svp* throughout the SOP lineage did not lead to a significant reduction in Or67d OSN number (Figure 6F). These results indicate that Svp functions downstream of Pnt to define Naa fate, but it is insufficient alone to confer Naa fate when ectopically expressed.

### The function of Pnt in Naa/Nab differentiation is likely to be independent of MAPK signal transduction

Pnt belongs to the ETS transcription factor family (Cooper et al., 2014; Klambt, 1993). Studies of both *Drosophila* and vertebrate homologs (ETS-1, ETS-2, Elk-1) have demonstrated that these transcription factors are MAP kinase-regulated nuclear effectors of the EGFR, JNK and p38 signaling pathways in diverse developmental and tissue contexts (Wasylyk et al., 1998; Whitmarsh et al., 1997). To determine whether Pnt is regulated in a similar manner in the at1 lineage, we performed RNAi on several ligands, receptors and cytoplasmic components of each of these signal transduction cascades. Although the size of the antenna and number of OSNs was altered in some cases (potentially reflecting developmental roles for these signaling pathways in this organ), we never observed doublets of Or67d OSNs (Figure S4C). We extended this analysis by simultaneous RNAi of components from two of these signaling cascades, but again failed to observe a pnt-like phenotype in Or67d OSN specification (Figure S4C). These results suggest that upstream regulation of Pnt in the at1 SOP lineage occurs in a different way to its other known developmental roles, potentially through regulation of its expression in distinct cells within this lineage (Figure 6B).

### A universal requirement for Pnt in OSN lineages

The at1 lineage is unique because it is the only sensillar subtype that harbors a single OSN (of Nab fate). All other olfactory sensilla contain two (Nab/Nba), three (Naa/Nab/Nba) or a maximum of four (Naa/Nab/Nba/Nbb) OSNs (Figure 7A). The at1 Naa cell that adopts Nab fate in the absence of Pnt is, in wild-type flies, originally destined for apoptosis (Figure 7B). To test if Pnt has a broader function in SOP lineages, we examined the expression of receptors in other sensilla classes in control and *pnt* RNAi antennae by RNA *in situ* hybridization. In the four classes of two-OSN sensilla tested (ab2, ab3, ai2, ac3), duplication of the Nab-derived OSN was observed, while expression of receptors in the other (Nba) OSN was unchanged (Figure 7C-D and Figure S5A-B). A similar phenotype occurs in the four-OSN ab1 sensilla: in this case, the Naa cell appears to transform from one OSN fate (Or10a-expressing) to another (Or92a-expressing), while the other two OSNs (Nba/Or42b and Nbb/Gr21a) are unaffected (Figure 7E and Figure S5A-C). These results are consistent with a conserved role of Pnt in the SOP lineages giving rise to these different classes of sensilla.

**Figure 7.**
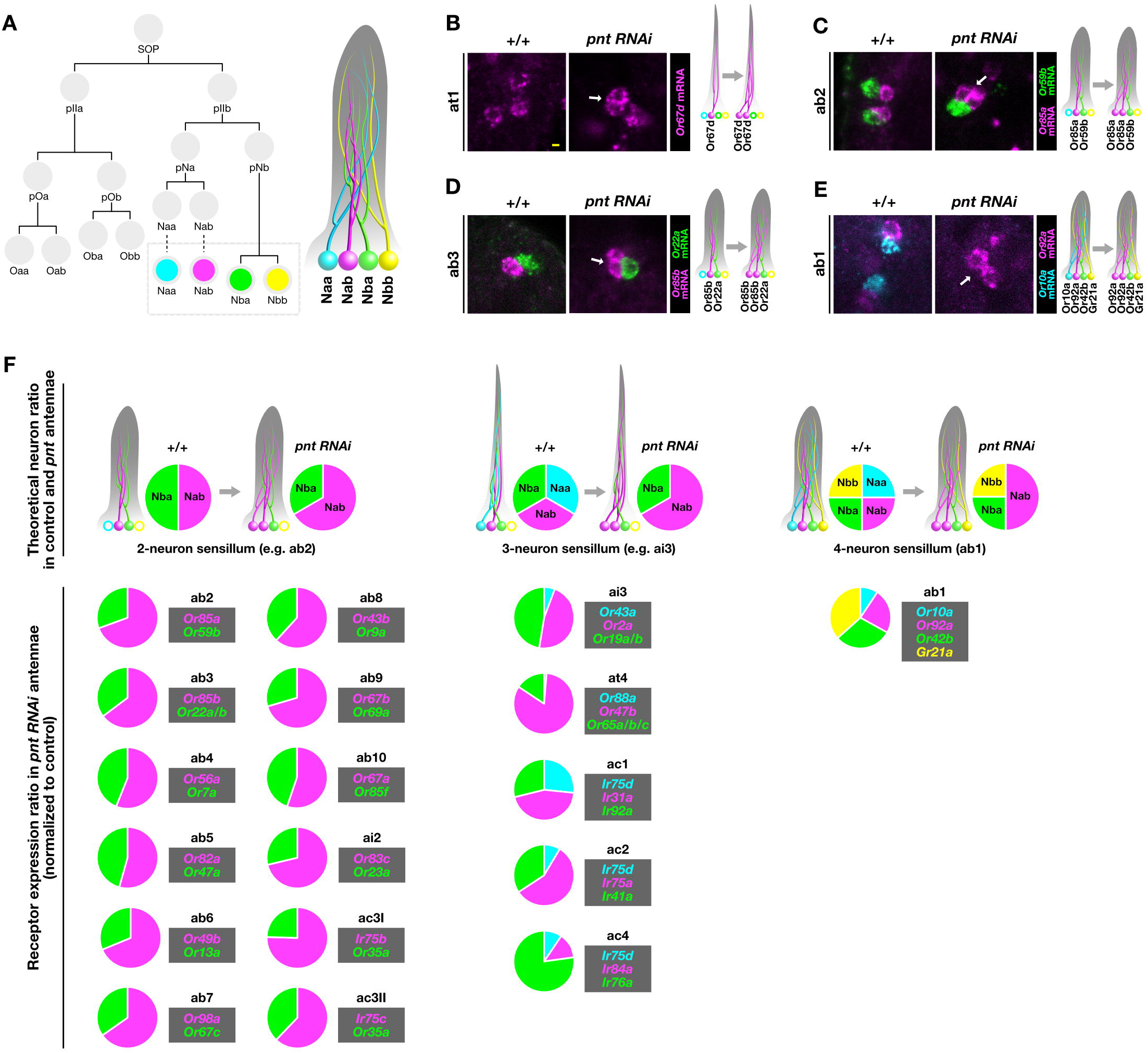
Loss of Pnt results in Naa-to-Nab transformations in diverse sensillar subtypes. (A) A sensillum can contain up to four OSNs through differentiation of Naa (cyan), Nab (magenta), Nba (green), Nbb (yellow) terminal daughter cells originating from a single SOP lineage. (B) Representative image of RNA FISH for *Or67d* (magenta) in at1 sensilla in control and *pnt* RNAi antennae. In *pnt* RNAi antennae Or67d expressing OSNs are duplicated in at1 sensilla (arrow). A schematic of the proposed Naa-to-Nab fate transformation is show on the right (color scheme as in (A)). Scale bar = 2 μm. The open circles in this and other schematics represent OSN precursors that have undergone apoptosis. (C) Representative image of RNA FISH for *Or85a* (magenta) and *Or59b* (green) in ab2 sensilla in control and *pnt* RNAi antennae. In *pnt* RNAi antennae, *Or85a* OSNs (Nab) are duplicated (arrow), while *Or59b* OSNs (Nba) are unaffected. (D) Representative image of RNA FISH for *Or85b* (magenta) and *Or22a* (green) in ab3 sensilla in control and *pnt* RNAi antennae. In *pnt* RNAi antennae, *Or85b* OSNs (Nab) are duplicated (arrow), while *Or22a* OSNs (Nba) are unaffected. (E) Representative image of RNA FISH for *Or92a* (magenta) and *Or10a* (cyan) in ab1 sensilla in control and *pnt* RNAi antennae. In *pnt* RNAi antennae, *Or92a* OSNs (Nab) are duplicated (arrow), while *Or10a* OSNs (Naa) are lost. (F) Top: theoretical ratios of OSN types in 2-, 3- and 4-neuron sensilla in control antennae, and in *pnt* RNAi antennae, assuming Naa-to-Nab fate transformation (i.e., loss of Naa OSNs, and increased in Nab OSNs). Bottom: experimentally-determined OSN ratios in all sensilla in *pnt* RNAi antennae using as a proxy the normalized ratios of olfactory receptor mRNA expression from antennal transcriptomes (see Figure S5F).

To expand this analysis to all antennal olfactory SOPs, we compared olfactory receptor gene expression levels in control and *pnt* RNAi antenna by RNA-sequencing. Loss of Pnt leads to an overall reduction in the number of antennal OSNs (Figure S5D) with a commensurate decrease in transcripts for most olfactory receptors (Figure S5E), suggesting that it has an earlier, general function in SOP specification in the antennal disc (consistent with its expression in SOPs (Figure 6A-B). Despite this global effect, we reasoned that cell fate transformations within individual SOP lineages would be recognizable as changes in the relative expression of receptor genes in individual sensilla (Figure 7F and Figure S5F).

Indeed, by normalizing the transcript read counts of the receptors to the theoretical ratios of their underlying OSNs (see Methods and Figure S5F), we observed that all (12/12) two-OSN sensilla classes had augmented expression of the receptor expressed in Nab relative to the receptor in Nba (Figure 7F). The change is most simply explained by a duplication of the Nab OSN, as already determined in a subset of sensilla (Figure 7B-E, and Figure S5A). Similarly, in most (4/5) three-OSN sensilla classes and the four OSN ab1 class, the increase of the Nab olfactory receptor expression relative to the Naa olfactory receptor expression is consistent with an Naa to Nab fate transformation (Figure 7F). An exception was observed in ac4, where *Ir84a* (Nab) expression is greatly diminished, potentially reflecting a role for Pnt in activation of this receptor gene. Globally, however, these transcriptomic data are consistent with a universal role for Pnt in distinguishing Naa from Nab in olfactory sensory lineages.

## Discussion

The correct structure and function of neural circuits depends on information encoded in the genome that is enacted during development. Our work was motivated by the desire to link the genesis of the peripheral olfactory system in *Drosophila* with the organization of the mature circuitry. By combining genetic resources that permit labeling of small subsets of precursor cells with methods for immortalization of expression patterns within defined temporal windows during development, we have generated a fate map of the “complete” olfactory system. Furthermore, we identified the first lineage-specific driver for an OSN population and, through a genome wide RNAi screen, discovered a novel switch-like genetic determinant of OSN lineages.

## An olfactory sensory fate map

A significant challenge of fate mapping using genetic methods is the dynamic nature of gene expression patterns, which is reflected in the observation that many transgenic reporters for developmental genes do not necessarily label the same cells – or even cells related by lineage – throughout development. This appears to be the case in the *Drosophila* peripheral olfactory system, as illustrated by an earlier endeavor to establish a fate map, where the unstable expression properties of several reporters led to the likely mis-mapping of OSNs within the antennal disc (Li et al., 2013).

Our immortalization approach has been instrumental in circumventing this limitation to establish a fate map of the precursors of all OSN classes within the developing antennal disc. Such a resource adds a novel developmental perspective on the *Drosophila* olfactory circuitry, complementing the maps of glomerular innervations (Couto et al., 2005; Fishilevich and Vosshall, 2005; Silbering et al., 2011) and PN projections to higher brain centers (Jefferis et al., 2007). This map reveals that while the concentric spatial organization is partially maintained from SOP birth to the sensilla distribution, no such order is noticeably preserved in the organization of the axon projections of OSNs in the antennal lobe. Indeed, the interleaved arcs of the Ato- and Amos-expressing SOPs produce sets of OSNs (the Ir and Or olfactory subsystems, respectively) that are spatially segregated within the primary olfactory center (Couto et al., 2005; Silbering et al., 2011). This observation, together with the lack of apparent clustering of glomeruli derived from SOPs born in the same arc, suggests that birthplace does not confer OSNs with axon guidance properties that reflect the topography of their origins.

Future identification and immortalization of additional enhancer-GAL4 drivers with restricted antennal disc expression should help improve the resolution of the fate map, and identify other sensilla lineage-specific markers. It is unclear, for example, whether different SOP classes that are born in the same arc are spatially segregated. This does not appear to be the case for at1 SOPs, which are dispersed throughout its arc, albeit in a more superficial layer than other SOPs. Such a distribution raises the question of the developmental mechanisms that act within a particular arc to specify up to 5-6 different SOPs types (e.g., Amos arc 7).

This map also provides a useful basis for investigating the evolutionary plasticity of the peripheral olfactory system. Comparative neuroanatomy of *D. melanogaster* and the olfactory specialist *D. sechellia* has identified two classes of OSNs/sensilla (OR22a/ab3 and IR75b/ac3I) that have substantially increased in number in the latter species, potentially related to its distinct olfactory ecology (Dekker et al., 2006; Prieto-Godino et al., 2017). As these increases are likely to reflect a higher number of the corresponding SOPs in the disc, our map may guide efforts to determine how specific SOP classes may change in abundance during evolutionary adaptation of the olfactory system.

## Transcriptional determination of OSN lineages

Previous genetic analyses have identified a number of developmental determinants that act at different time-points in OSN lineages (Barish and Volkan, 2015), including the proneural molecules Amos and Ato, which distinguish SOPs producing the Or and Ir lineages (Goulding et al., 2000; Gupta and Rodrigues, 1997; zur Lage et al., 2003) and iterated Notch-dependent nuclear signals to define asymmetries within many of the SOP lineage divisions (Endo et al., 2007; Endo et al., 2011). Comparatively little is known about the mechanisms by which different SOPs are specified within the antennal disc to give rise to the enormous diversity of different sensilla and OSN classes. The best-characterized early patterning determinant is Rn, which is required for the specification of a subset of OSNs (Li et al., 2015; Li et al., 2013). Our fate map is consistent with these phenotypes, as the OSN classes that are lost in *rn* mutants are precisely those that derive from Rn-positive arcs in the antennal disc.

Previous large-scale RNAi screening of transcriptional determinants of OSN fate focused on those that have late functions in olfactory receptor choice. These efforts identified a small set of factors that act in a combinatorial manner to activate or repress olfactory receptor expression in specific OSN classes (Alkhori et al., 2013; Bai et al., 2009; Jafari et al., 2012). Such factors are likely to act only at the end of more elaborate gene regulatory networks that ensure the specification of SOP type, and determination and coordination of OSN receptor expression and axon targeting (Hueston et al., 2016; Tichy et al., 2008). The genome-wide, “constitutive” RNAi screen of transcriptional regulators performed here has identified a large number of new molecules that are likely to function in several of these processes. Future analysis of their expression and precise phenotypes using the at1 driver should enable determination of their function within this and other OSN lineages.

## A novel switch-like determinant in OSN lineages

We focused here on the role of the ETS homolog, Pnt, because of its unique mutant phenotype which reveals a role in limiting, rather than determining, Or67d neuron specification. With our at1 lineage marker, protein expression and gain- and loss-of-function analyses, we provide evidence that this transcription factor has a switch-like function in distinguishing the terminal Svp-expressing Naa cell from its Svp-negative sibling Nab. Interestingly, this role of Pnt is distinct from other roles of this transcriptional factor where it serves as a nuclear read-out of various MAPK signaling pathways, as none of these appear to be important in this context.

Our antennal transcriptomic analysis indicates that this role of Pointed is likely to be universal in olfactory sensilla. The majority of these contain two OSNs, reflecting the role of Pnt (and Svp) in diverting the fate of Naa cells away from neural differentiation and toward apoptosis. Despite the potential for each sensillum to specify four OSNs, this is only realized in the ab1 sensillum class. It is unclear why so many are removed by programmed cell death. One possibility is that the higher numbers of neurons compartmentalized within a sensillum would result in substantial non-synaptic electrical interference between OSNs (Su et al., 2012), with potential detrimental consequences for odor coding, especially if OSNs have similar tuning properties (Vermeulen and Rospars, 2004).

The switch-like function is not the only role of Pnt in the antenna, as it also contributes to the specification of the correct global number of SOPs, and may more directly regulate the expression of specific olfactory receptor genes (e.g., *Ir84a).* Moreover, Pnt’s broad expression in the non-neuronal sublineage suggests it may also participate in support cell development. Consistent with this possibility, preliminary observations of *pnt* RNAi antennae reveal abnormalities in the morphology of the sensillar cuticular hairs (e.g., Figure 5E and S5D), which are determined, in part, by support cells (Keil, 1999).

## Linking developmental-omics and *in situ* imaging

Recent advances in genome-editing and single-cell RNA-seq or chromatin profiling technologies have been revolutionary for documenting and classifying cell-type diversity in the nervous system (and other tissues), as well as in defining their lineage relationships (e.g., (Raj et al., 2018; Zhong et al., 2018)). With our OSN lineage driver, we may now exploit such technologies to examine the gene expression and epigenetic states of the at1 lineage from birth to maturity, and how these may be influenced by internal state and environmental conditions, both of which have been shown to impact OSN specification (Jafari and Alenius, 2015). While cellular-resolution level transcriptomic/epigenomic data is undeniably important to understand neural development, the combination of these with tools for visualization of specific lineages *in vivo* is essential for a complete view of how structural and functional diversity develops in the nervous system.

## Acknowledgements

We are grateful to Mattias Alenius, Thomas Auer, Konrad Basler, Hugo Bellen, Yu Cai, Marco Cantoni, Christian Fankhauser, Veit Grabe, Fisun Hamaratoglu, Yasushi Hiromi, Michael Hoch, Yuh-Nung Jan, Andrew Jarman, Kaan Mika, Marco Milán, Pavan Ramdya, Silke Sachse, Pelin Volkan, Leslie Vosshall, Ryohei Yagi, Petra zur Lage, the Bloomington *Drosophila* Stock Center (NIH P40OD018537), the Vienna Drosophila Resource Center, the KYOTO Stock Center (Kyoto DGGR of Kyoto Institute of Technology), the Developmental Studies Hybridoma Bank (NICHD of the NIH, University of Iowa), and the Lausanne Genomic Technologies Facility for their generous sharing of reagents and resources. We thank Scott Barish, Sebastian Cachero, Yu Cai, Erika Dona, Pelin Volkan and members of the Benton laboratory for discussions and comments on the manuscript. Research in R.B.’s laboratory is supported by the University of Lausanne, an ERC Consolidator Grant (615094) and the Swiss National Science Foundation (31003A_166646).

## Author Contributions

P.C.C. conceived the project, designed, performed and analyzed most experiments. S.C. performed RNA *in situ* hybridization and prepared samples for RNA-seq. L.W. performed statistical analyses of RNA-seq data. R.B. supervised the project. P.C.C. and R.B. wrote the paper, with input from other authors.

## Methods

### *Drosophila* stocks

*Drosophila* stocks were maintained on standard cornmeal medium under a 12:12 h light/dark cycle at 25°C, unless otherwise stated. The mutant and transgenic lines used are listed in Table S2.

## GAL4 driver immortalization

Embryos of flies arising from the desired genotypic cross were collected over 8 h and allowed to develop at 19°C until the larvae began pupation (typically after ~7 days under our culture conditions). All non-white pupae (representing older ages) were discarded, while the other larvae and white pupae (i.e., aged ~0-8 h BPF) were subjected to 24 h at 29°C to heat inactivate the GAL80^ts^. Animals were returned to 19°C to continue development until eclosion after a further ~6-7 days. For late immortalization of *en-GAL4,* heat-inactivation was carried out on pupae aged ~9-15 h APF.

For at1 lineage analysis in the adult, embryos were collected over a period of either 9 h or 15 h (representing overday and overnight collections, respectively) at 19°C, and aged until the larvae began pupation. All the older pupae were discarded such that the remaining larvae/pupae (aged between ~0-9 h APF or ~0-15 h APF for the different collections) were placed at 29°C for 24 h to heat-inactivate the GAL80^ts^ (the stability of at1 driver expression allows the application of a wider age window than for other GAL4 drivers). The pupae were returned to 19°C to continue development until they were ready to be assayed.

The analysis of at1 lineage development in the pupae was performed as above, but to ensure tight synchronization of development, only the white pupae (aged ~0 h APF) were collected. These were subjected up to a maximum of 24 h heat-inactivation of GAL80^ts^ at 29°C, before they were returned to 19°C for development. Pupal antennae were dissected at specific time points between 18 h APF to 27 h APF (i.e., within or after the heat-inactivation window).

## RNA interference

To maximize the efficiency of RNAi, larvae of the desired genotype were raised at 27°C (which enhances GAL4 induction) from 24 h after egg-laying until the animals were ready to be analyzed. For the minority of transgenic RNAi lines that resulted in pupal lethality, larvae were allowed to develop at 25°C.

## Clonal generation

Heat-shock clones and MARCM clones were generated according to established methods (Ito et al., 1997; Lee and Luo, 1999). Embryos were collected over 15 h and raised at 25°C until 48-63 h after larval hatching. These larvae were subjected to 30 min heat shock in a 37°C water bath and returned to 25°C to continue development. For the examination of antennal SOP lineage development, white pupae were collected to re-synchronize their developmental timing and aged until 20-24 h APF.

## Histology and immunocytochemistry

Immunofluorescence on adult brains was performed according to a standard protocol (Sanchez-Alcaniz et al., 2017). Immunofluorescence and RNA fluorescence *in situ* hybridization (FISH) on whole-mount adult antennae were performed as described (Saina and Benton, 2013) with slight modifications: (i) antennae were fixed for 3 h at 4°C, (ii) RNA probes were denatured at 80°C for 10 min and (iii) incubation with anti-Digoxigenin (DIG)-POD or anti-Fluorescein (FITC)-POD lasted for 36 h.

For immunofluorescence on larval antennal discs and pupal antennae, animals were placed in phosphate buffered saline (PBS) and severed below the head to expose the discs or antennae before fixation in 4% paraformaldehyde in PBS + 0.2% Triton X-100 (PBT) for 1.5 h at 4°C. Following ~5-10 quick rinses with PBT, excess tissues were removed. Samples were blocked in 5% normal goat serum diluted in PBT for 1 h at room temperature (RT) before incubation in primary antibodies (diluted in blocking solution) for 48 h at 4°C. After five rounds of washes with PBT over 3 h, secondary antibodies (diluted in blocking solution) were added and left for 48 h at 4°C. Following another five rounds of washes with PBT over 3 h, samples were equilibrated overnight at 4°C in Vectashield mounting medium. During mounting onto a bridged slide, all unwanted tissues and head structures were cleaned from the antennal discs and pupal antennae.

The antibodies used in this work are listed in Table S3). All microscopy was performed using a Zeiss LSM 710 laser scanning confocal microscope. The raw confocal images were processed (cropping, brightness/contrast adjustment, and color channel separation) in Fiji (Schindelin et al., 2012).

## Molecular biology

The construction of plasmid templates and synthesis of DIG- and FITC-labeled RNA probes was performed as previously described (Benton et al., 2007; Saina and Benton, 2013). The primers used to amplify the desired cDNA or genomic fragments (or previously-described probes) are listed in Table S4.

## Total RNA extraction from antennae

Antennal RNA was extracted from four biological replicates of *pnt* RNAi flies (constitutive driver crossed to *UAS-pnt^RNAi-KK100473^*GAL4 driver immortalization) and their paired controls (constitutive driver crossed to the same strain without the RNAi transgene: VDRC control 60100). For each pair of biological replicates, animals were grown under identical conditions and RNA was extracted in parallel.

For each biological replicate, the third antennal segments from 500-1000 flies were harvested via snap-freezing in a mini-sieve (Gomez-Diaz et al., 2013). Antennae were transferred to a 1.5 ml Eppendorf tube on ice and homogenized manually with a tissue grinder. Total RNA was extracted from the homogenized antennae using a standard TRIzol/chloroform protocol. Briefly, TRIzol reagent was added to a final volume of 1 ml and the tube shaken vigorously manually for 1 min. After chilling on ice for 5 min and centrifugation (13,000 rpm for 15 min at 4°C, for all subsequent centrifugation) the supernatant was transferred to a new tube. 200 μl of chloroform was added, and the tube shaken vigorously for 15 s and left at RT for 3 min, before re-centrifugation. The upper aqueous phase was transferred to a new tube, and RNA was precipitated with an equal volume of prechilled isopropanol. The mixture was incubated for 10 min at RT and centrifuged for 15 min. The resultant RNA pellet was washed with 1 ml prechilled 75% ethanol, air-dried, and reconstituted in 20 μl water.

## RNA library preparation and sequencing

RNA quality was assessed on a Fragment Analyzer (Advanced Analytical Technologies, Inc., Ankeny, IA, USA) and all RNAs had an RQN between 7.9 and 10. From 100 ng total RNA, mRNA was isolated with the NEBNext Poly(A) mRNA Magnetic Isolation Module. RNA-seq libraries were then prepared from the mRNA using the NEBNext Ultra II Directional RNA Library Prep Kit for Illumina (New England Biolabs, Massachusetts, USA). Cluster generation was performed with the resulting libraries using the Illumina TruSeq SR Cluster Kit v4 reagents and sequenced on the Illumina HiSeq 2500 using TruSeq SBS Kit v4 reagents (Illumina; San Diego, California, USA). Sequencing data were demultiplexed using the bcl2fastq Conversion Software (version 2.20, Illumina; San Diego, California, USA).

## RNA-seq data analysis pipeline

Purity-filtered reads were adapters and quality trimmed with Cutadapt (version 1.8, (Martin, 2011)). Reads matching to ribosomal RNA sequences were removed with fastq_screen (version 0.11.1). Remaining reads were further filtered for low complexity with reaper (version 15-065, (Davis et al., 2013)). Reads were aligned to the *Drosophila melanogaster* BDGP6.86 transcriptome using STAR (version 2.5.3a, (Dobin et al., 2013)) and the estimation of isoform and gene abundance was computed using RSEM (version 1.2.31,(Li and Dewey, 2011)). Gene-level estimated counts from RSEM were used as input for the statistical analysis. Gene-level TPM (transcripts per kilobase million) from RSEM were used for the calculations of read count ratios between genes expressed in a sensilla type.

Statistical analysis was carried out using R (version 3.4.4). The *pnt* RNAi antennae were compared to controls with a paired samples design. The RSEM gene-level data was read in with the Bioconductor package tximport (version 1.6.0, (Soneson et al., 2015)). Fold changes and *p*-values were calculated with the R Bioconductor package DESeq2 (version 1.18.1), which uses negative binomial GLM fitting and Wald statistics (Love et al., 2014). Genes with zero estimated counts in all samples were removed prior to normalization and model fitting, leaving 14233 genes in the analysis. Default settings were used for the estimateSizeFactors() and estimateDispersions() functions. For multiple testing correction, the *p*-values were adjusted together by the Benjamini-Hochberg method, which controls the false discovery rate (Benjamini and Hochberg, 1995). The independent filtering option for *p*-value adjustment was turned off.

RNA-seq data will be deposited in GEO upon publication.

## Supplementary Information

**Figure S1.**
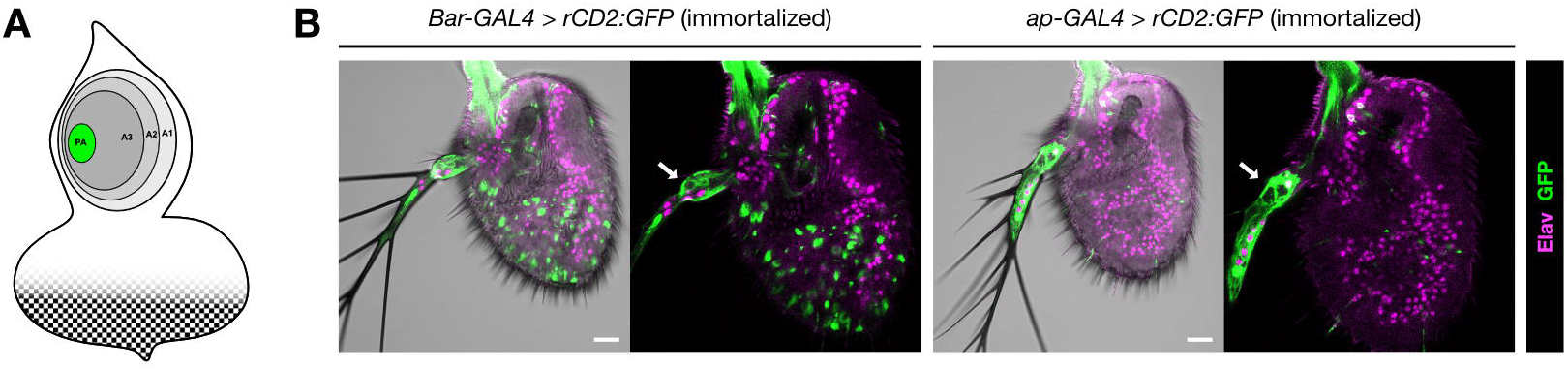
Arista neurons originate from SOPs located at the center of the concentric rings. (*related to Figure 1*) (A) The arista SOPs are located in the presumptive arista (PA) area (green) of the antennal disc. (B) Immortalization of *Bar*-GAL4 and *ap*-GAL4 in a 4 h BPF-20 h APF window produced GFP labeled neurons (green) in aristal neurons (GFP signal elsewhere in the antenna is non-neuronal background). Scale bar = 20 μm. The genotypes of the flies used in the figures are listed in Table S1.

**Figure S2.**
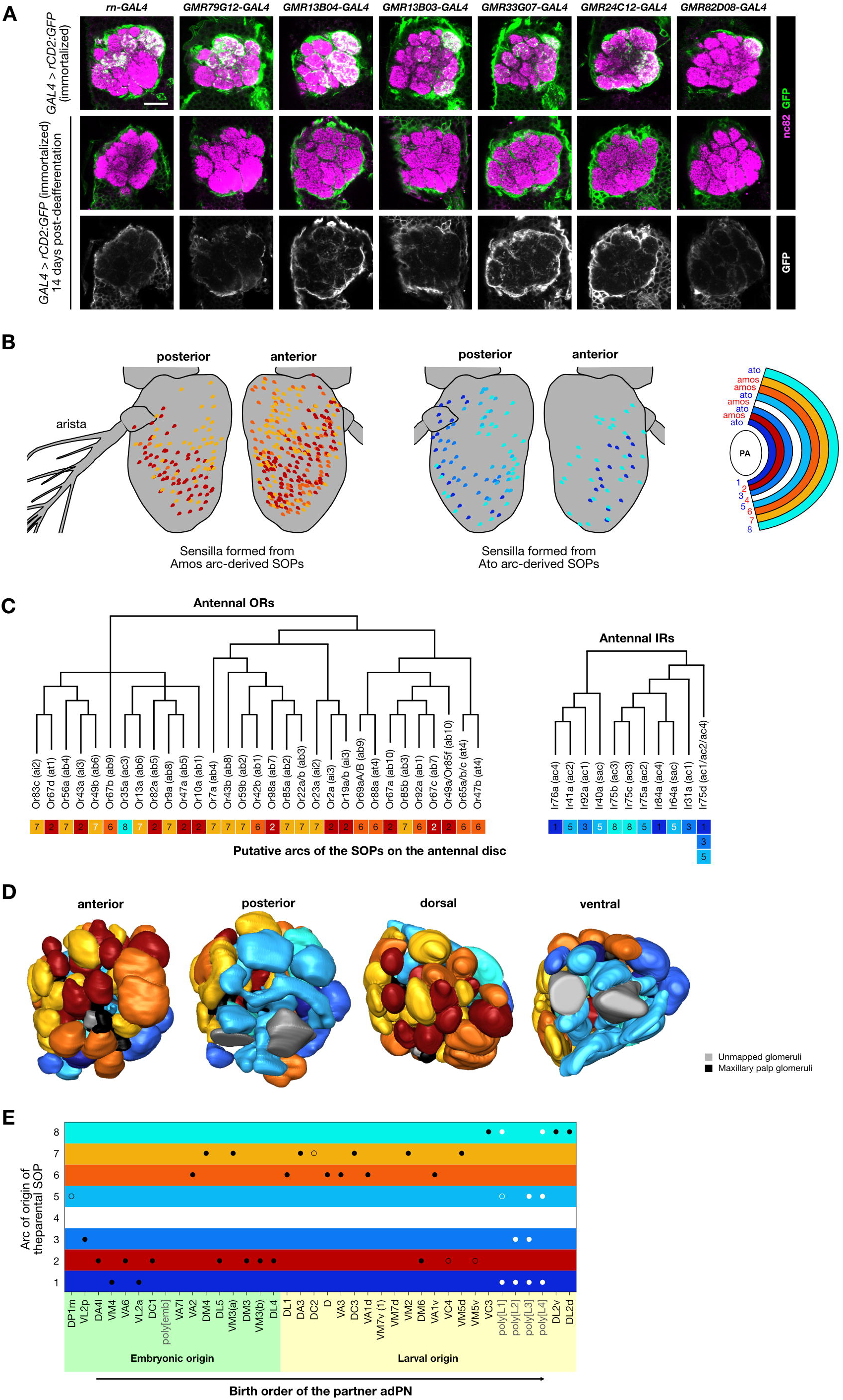
Correlations between the olfactory fate map and their neuroanatomical properties. (*related to Figure 2*) (A) Row 1: immortalized *enhancer*-GAL4 driven expression of rCD2:GFP (α-GFP; green) in OSN axons innervating different subsets of glomeruli labeled by nc82 (magenta). Rows 2-3: immortalized *enhancer*-GAL4 driven expression of rCD2:GFP is lost in the antennal lobe glomeruli 14 days after the antennae were surgically removed. Scale bar = 20 μm. (B) Distribution of different sensilla classes (from (Grabe et al., 2016)), color-coded for their antennal disc arc of origin (one color may represent many different sensilla types that arise from the same arc). Sensilla originating from both Amos-positive arcs (red-yellow gradient) and Ato-positive arcs (blue-cyan gradient) are show separately for clarity; both partially preserve their relative positions in the antenna: the internal arcs produce sensilla that have lateral (L) distributions while the peripheral arcs produce sensilla with more medial (M) distributions. (C) The relationship between Or/Ir protein phylogeny and the arc of origin of the corresponding SOPs (derived from the sensilla fate map in Figure 2F). The Or phylogenetic tree is from (McBride and Arguello, 2007); the Ir phylogenetic tree is from (Silbering et al., 2011). The arcs from which the OSNs originate are color-coded as in (B). Arc labels in black: high confidence OSN/sensillum mapping; arc labels in white: ambiguous OSN/sensillum mapping. (D) Different 3D surface views of the right antennal lobe (from (Grabe et al., 2014)), in which glomeruli are colored (as in (B)) according to the arc of origin of the SOPs from which the corresponding OSNs arise. (E) Relationship between OSN birthplace and PN birth order. OSN classes (defined by their glomerular innervation) are positioned on the vertical axis according to the arc of origin of the SOPs from which they arise, and ordered on the horizontal axis by the birth timing of their synaptic partners produced in the anterodorsal PN (adPN) neuroblast lineage (based on data from (Yu et al., 2010)). Black/white dots indicates mono-/polyglomerular OSNs, respectively (filled dots: OSNs from sensilla mapped with high certainty; unfilled dots: OSNs from ambiguously mapped sensilla).

**Figure S3.**
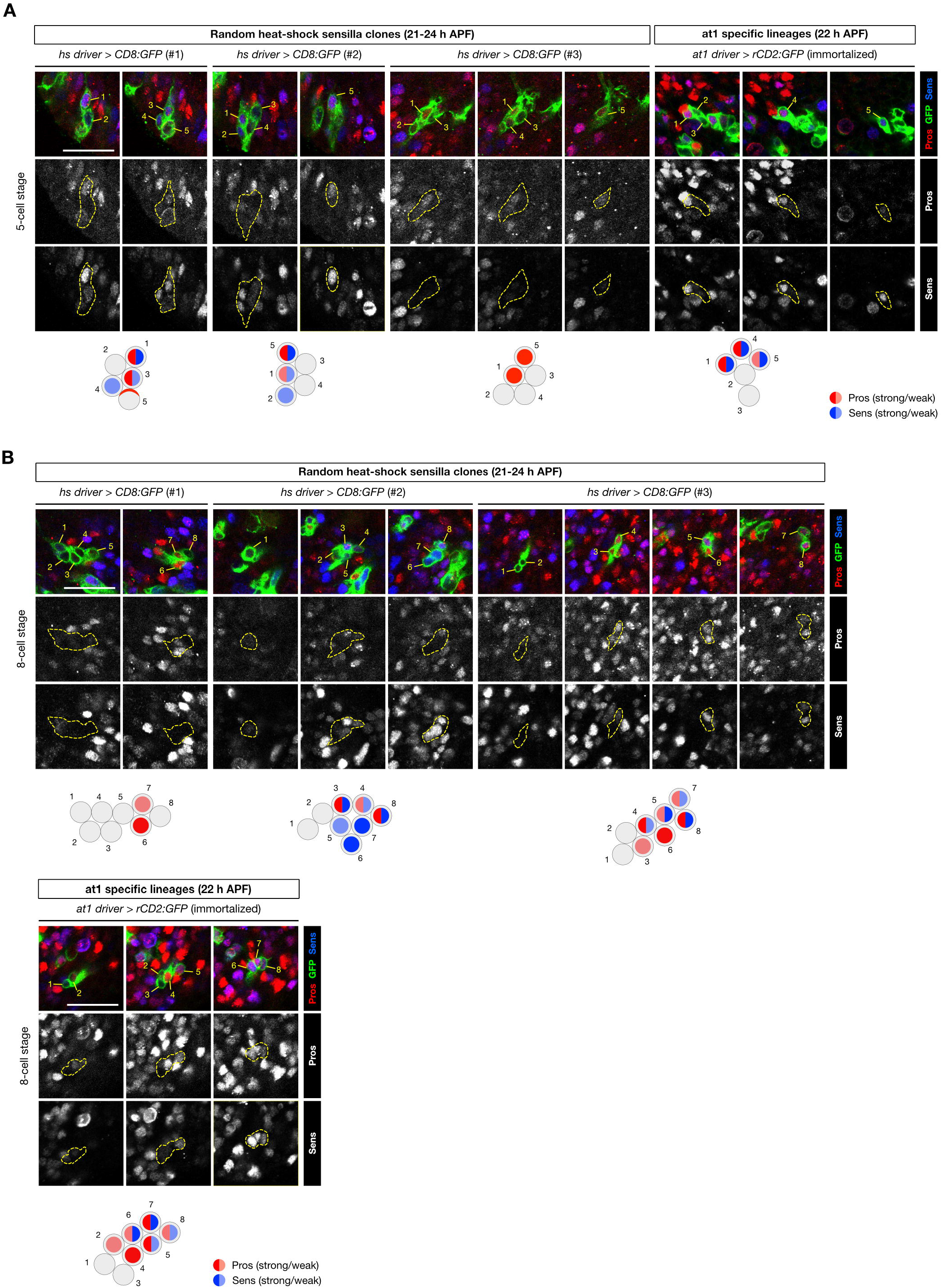
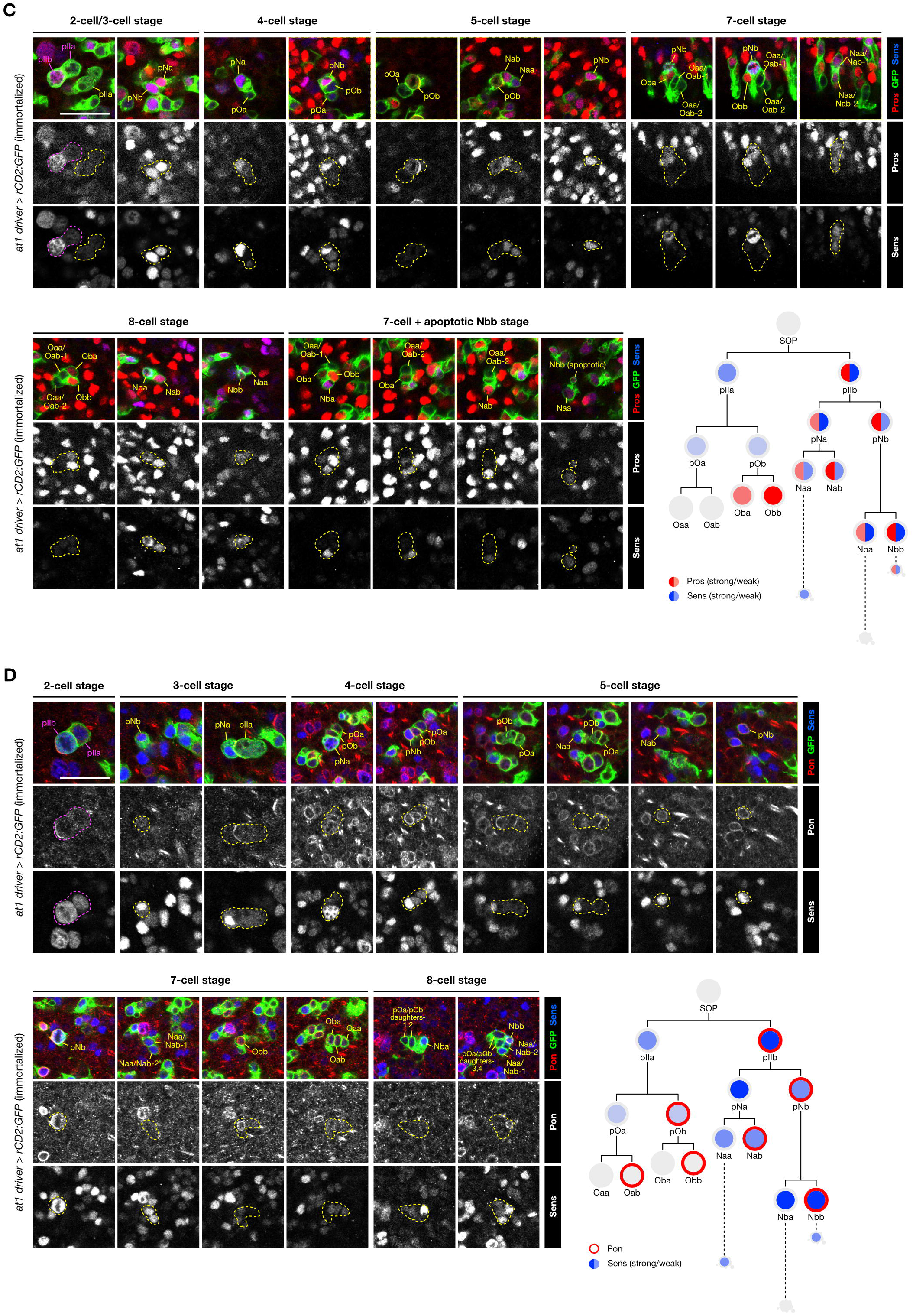
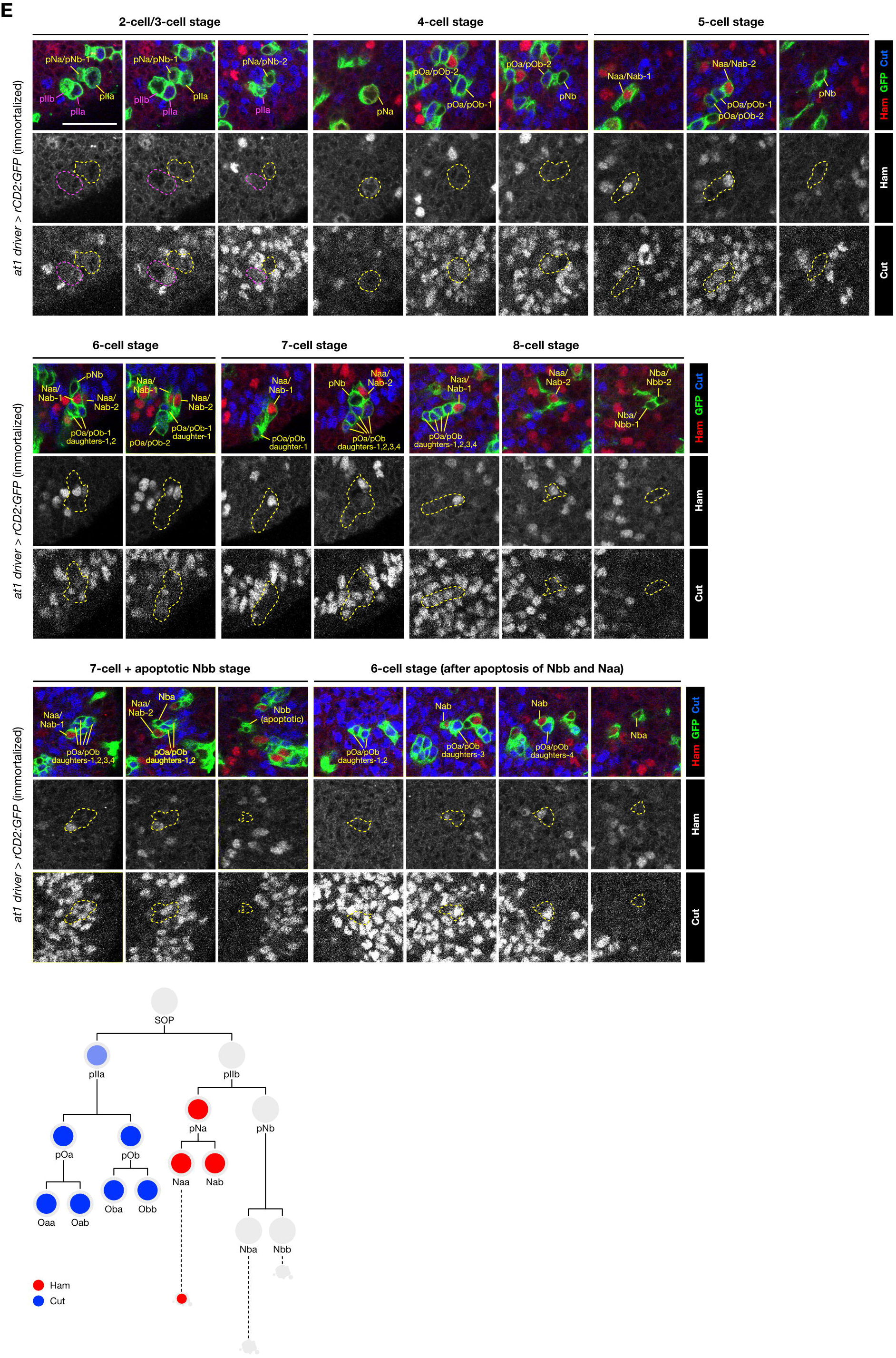
at1 lineages show invariant molecular marker expression. (*related to Figure 4*) (A) Left: the 5-cell stage of random SOP lineages (visualized by heat shock-based clonal GFP labeling (green); clonal generation time: 48-64 h after larval hatching)) in the developing pupal antennae at 21-24 h APF have different expression profiles for Pros (red) and Sens (blue). Right: by contrast, a 5-cell stage of the at1 lineage (visualized with the immortalized at1 driver) has highly predictable Pros and Sens expression profile. Schematic summaries of the expression patterns are shown below the images. Scale bar = 20 μm in this and other panels. (B) Top: the 8-cell stage of random SOP lineages (visualized by heat shock-based clonal GFP labeling as in (A)) in 21-24 h APF antennae have highly variable Pros (red) and Sens (blue) expression patterns. Bottom: by contrast, the 8-cell stage of the at1 lineage shows invariant Pros and Sens expression profiles. (C)-(E) Expression profiles of (C) Pros (red), Sens (blue), (D) Pon (red), Sens (blue), and (E) Ham (red), Cut (blue), in at1 lineages visualized with the immortalized at1 driver (green). Schematics summarizing the expression profiles are shown after each image series.

**Figure S4.**
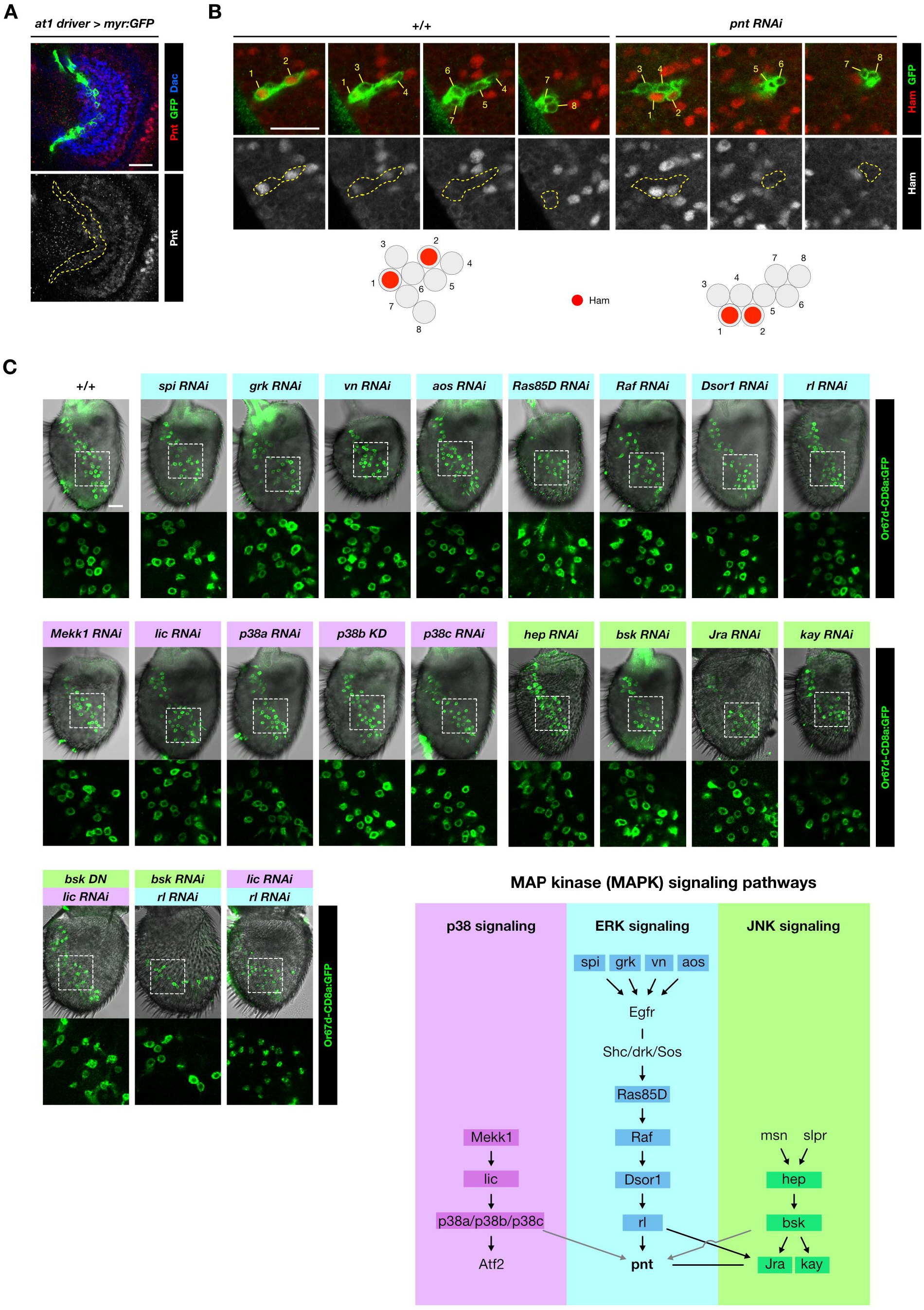
The role of Pnt in Naa fate specification is unrelated to MAP kinase signaling pathways. (*related to Figure 6*) (A) Pnt (red) is expressed at low levels in the at1 SOPs (green) of a 2 h APF antennal disc; Dac (blue) demarcates the A3 region. Scale bar = 20 μm in this and other panels. (B) Left: representative control 8-cell heat-shock clone (green) revealing two cells expressing Ham (red). Right: in a representative 8-cell *pnt* RNAi clone (green), Ham expression is unaffected. Schematics summarizing Ham expression in these clones are shown below the images. (C) Antennae expressing the Or67d-CD8a:GFP reporter in control animals or with single or double RNAi (using the constitutive driver) of genes in three main MAPK signaling pathways. For two experiments, non-RNAi-based tools were used: *bsk* dominant negative (DN) and *p38b* KD (kinase dead). The areas within the dashed boxes are magnified to show the cell bodies of GFP-expressing neurons; no doublets are observed, in contrast to *pnt* RNAi antennae (Figure 5E). Bottom right: schematic of the core components of p38 (purple), ERK (cyan) and JNK (green) signaling pathways. The black arrows/lines indicate known interactions in *Drosophila*, while the grey arrows indicate interactions reported for their mammalian homologs.

**Figure S5.**
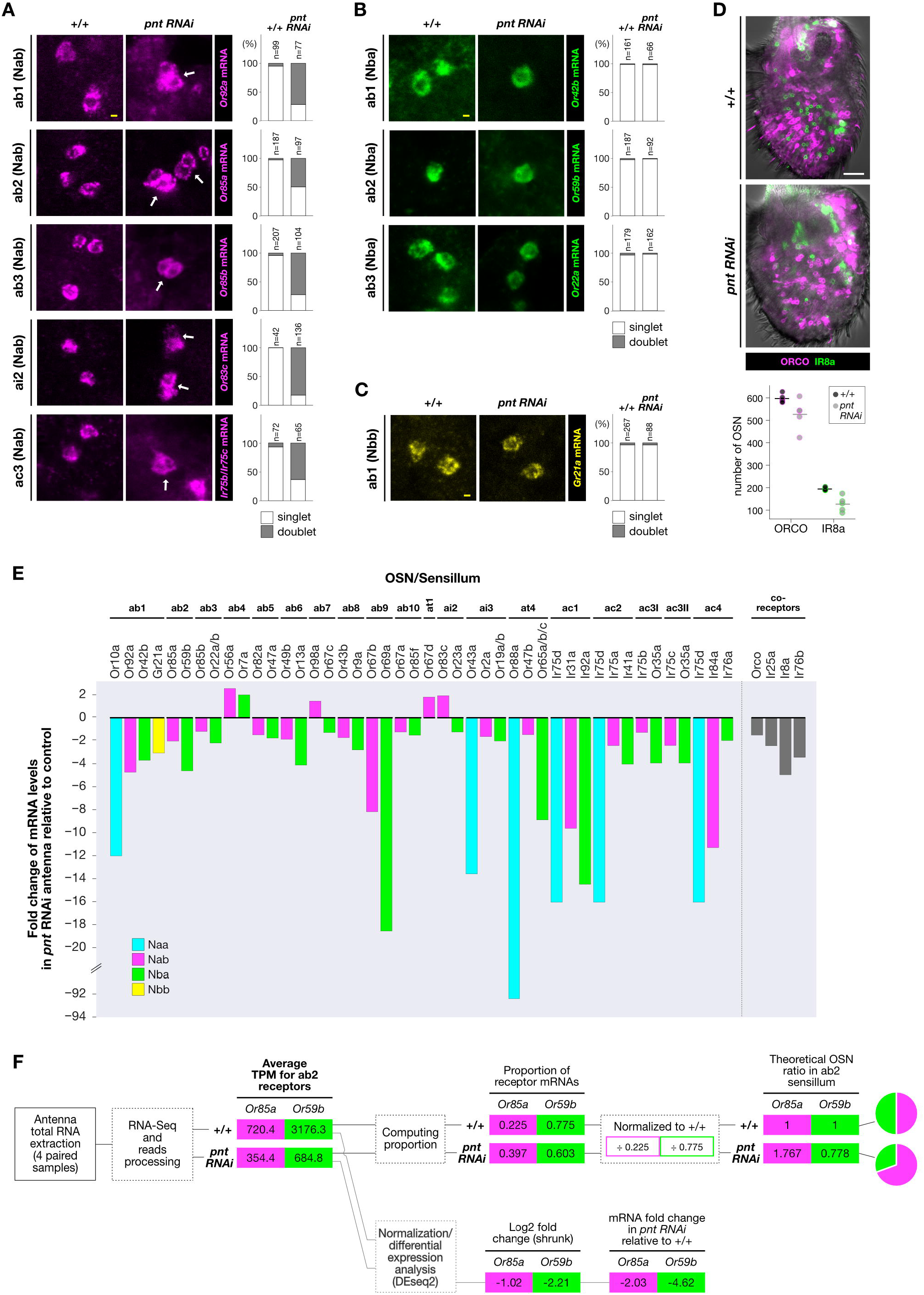
Loss of Pnt results in Naa-to-Nab transformations in diverse sensillar subtypes, and a global reduction in sensilla number. (*related to Figure 7*) (A) OSNs of Nab fate, as labeled by FISH probes against their olfactory receptor mRNAs (magenta), in the indicated sensilla classes in control and *pnt* RNAi antennae. *pnt* RNAi antennae exhibit doublets of Nab, although phenotypic penetrance was variable between different sensillar subtypes, possibly due to the contribution of other neural fate determinants. Yellow scale bar = 2 μm in this and other panels. (B) OSNs of Nba fate, as labeled by FISH probes against their olfactory receptor mRNAs (green), in the indicated sensilla classes in control and *pnt* RNAi antennae. (C) The OSN of Naa fate in ab1 sensilla, as labeled by a FISH probe against *Gr21a* mRNA (yellow), in control and *pnt* RNAi antennae. (D) Immunolabeling with α-Orco (magenta) and α-Ir8a (green) (which together label most antennal OSNs) in control and *pnt* RNAi antennae. The number of both Orco- and Ir8a-expressing OSNs is reduced upon loss of *pnt:* the distribution of the OSN counts is presented as a strip-plot (horizontal line denotes the sample mean). (n=5; Mann-Whitney U test: p=0.047 and p=0.006 for Orco- and Ir8a-expressing OSNs, respectively, comparing control and *pnt* RNAi genotypes). White scale bar = 20 μm. (E) Expression of olfactory receptors in *pnt* RNAi antennae relative to control antennae, based on four paired biological replicates. The expression of all antennal olfactory receptors in the *pnt* RNAi antennae are significantly different from their control counterparts (adjusted p<0.05), apart from *Or67c* (adjusted p=0.063) and *Or85b* (adjusted p=0.121). Statistics were computed with DESeq2. *p*-values were adjusted by the Benjamini-Hochberg method across all genes with non-zero counts. (F) Workflow to compute fold-change of mRNA levels (Figure S5E) and infer OSN numbers in sensilla in *pnt* RNAi antennae (Figure 7F) using, as an example, the read counts for ab2 receptors (*Or85a* and *Or59b).* TPM: transcripts per million.

**Table S1:**
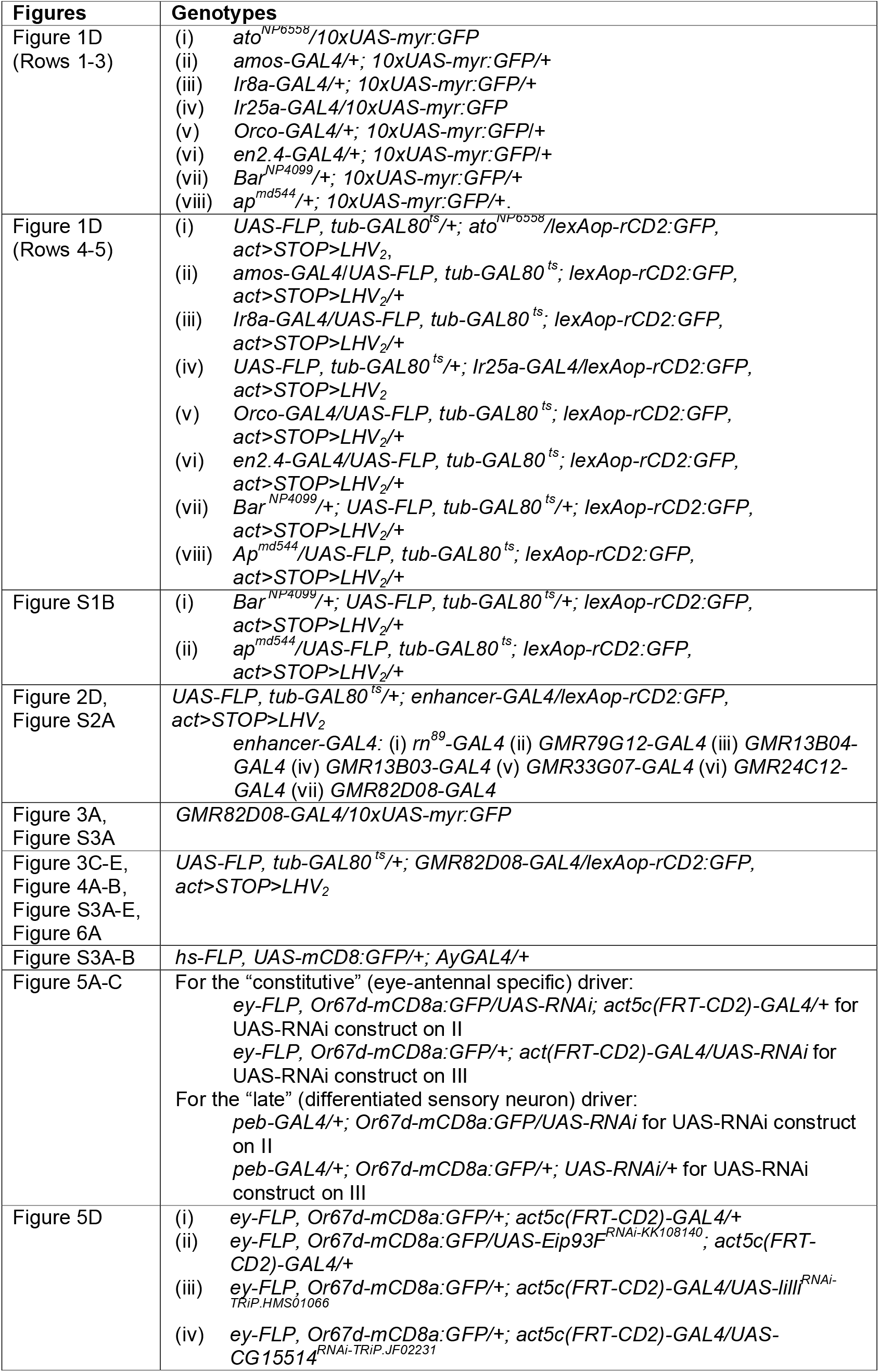
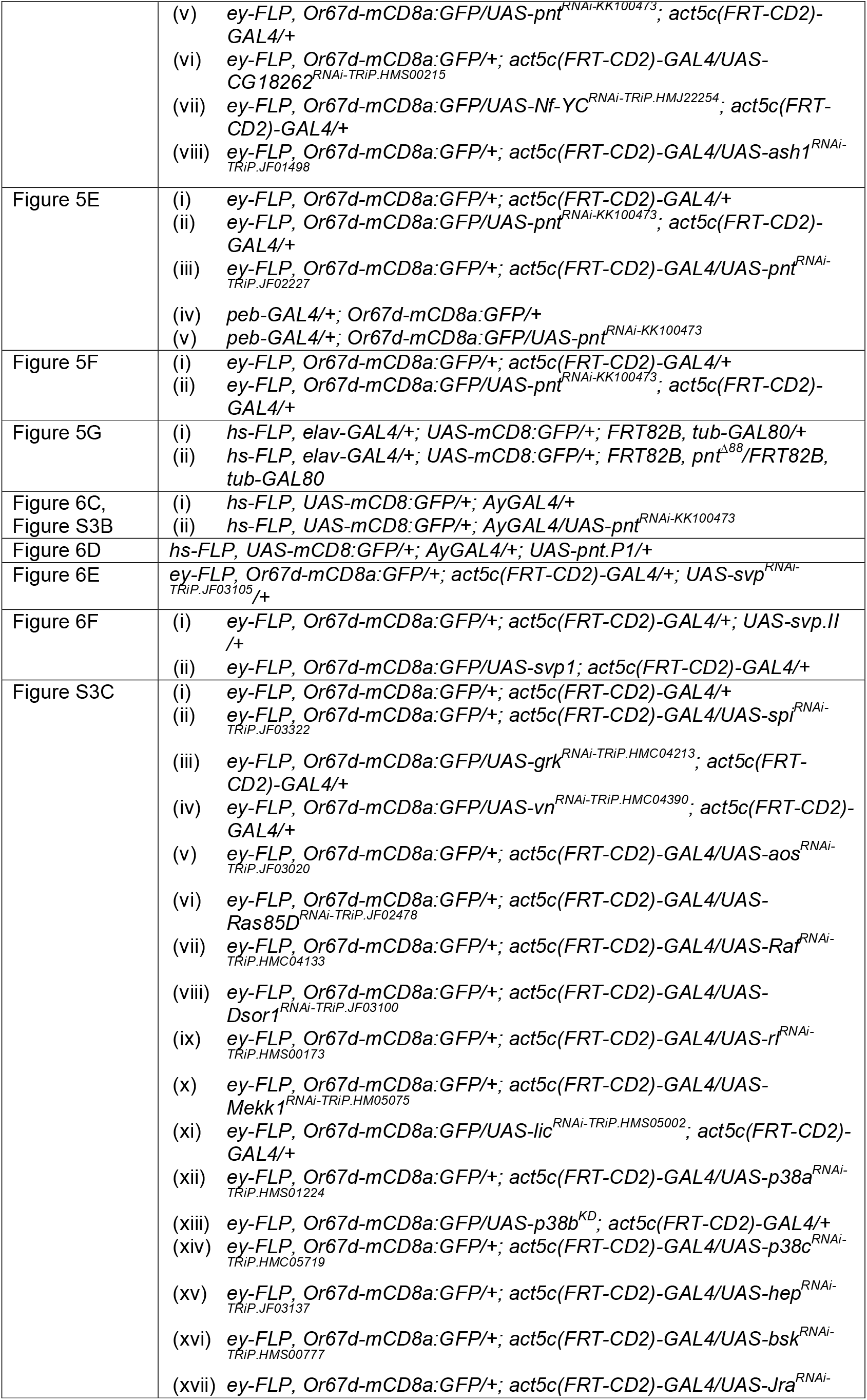
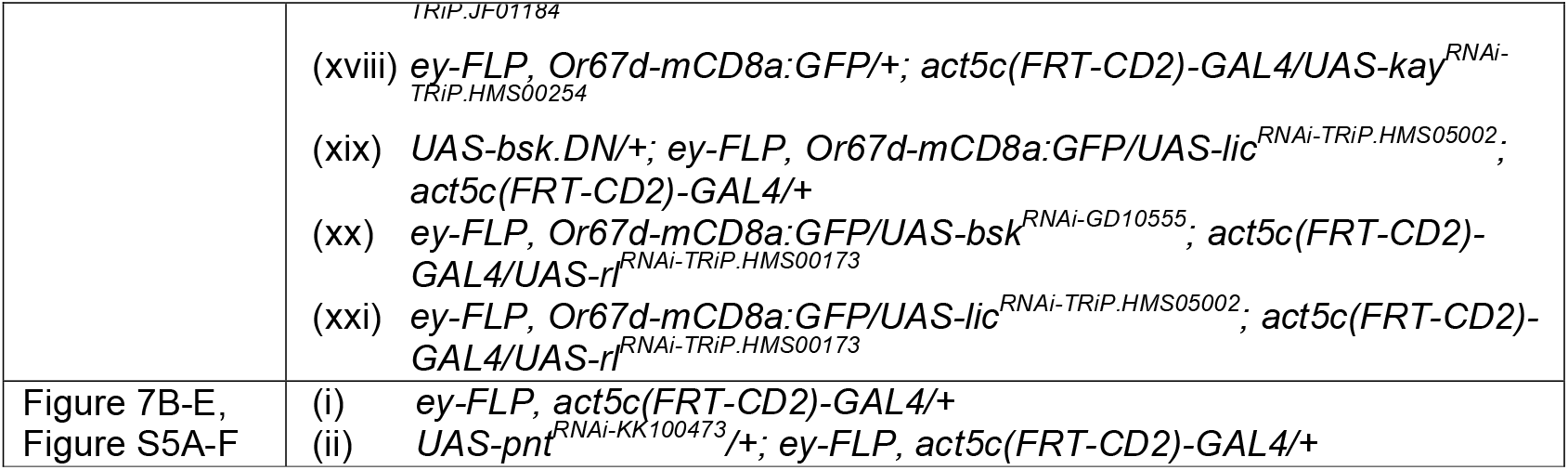
Genotypes of the experimental animals.

**Table S2.**
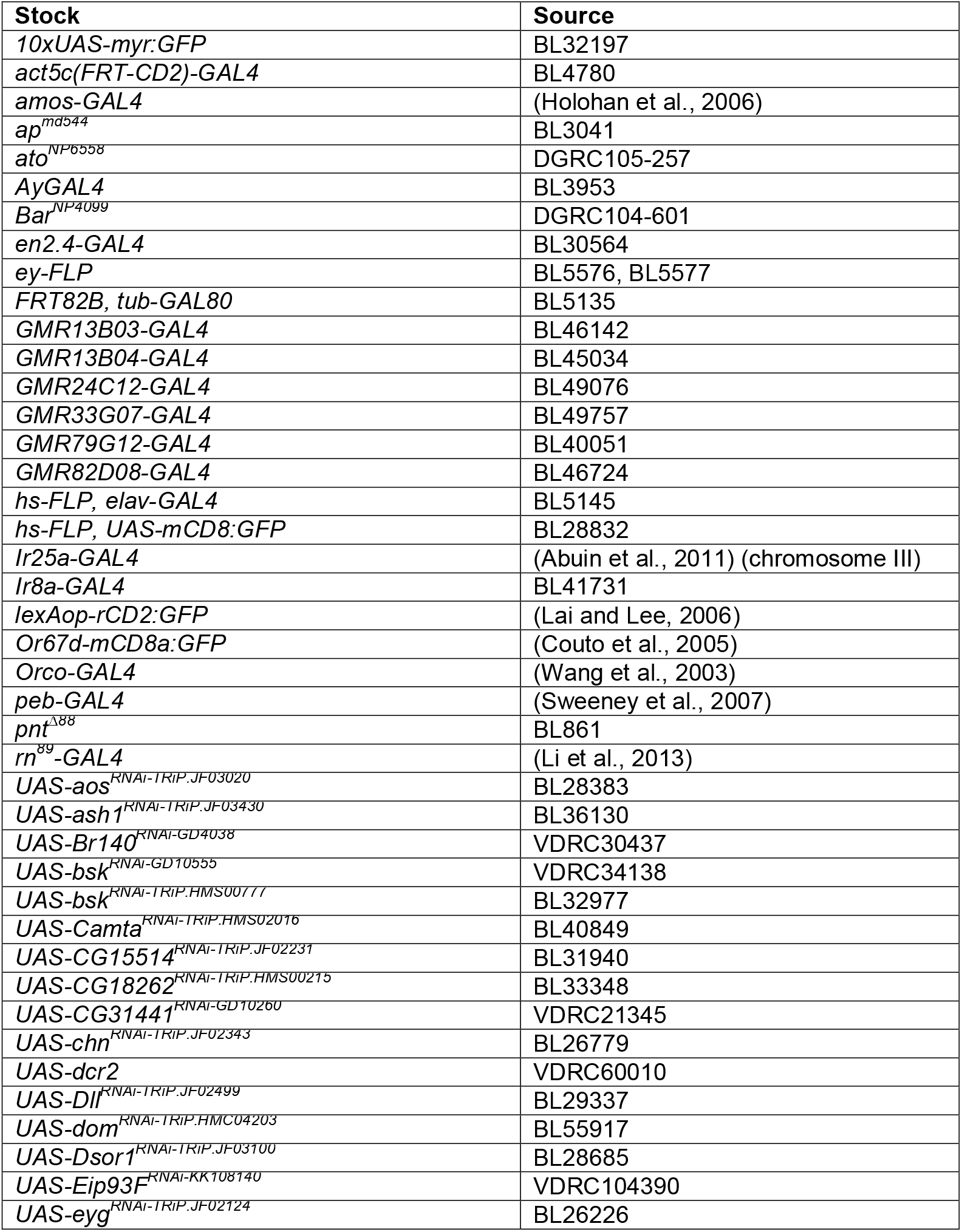
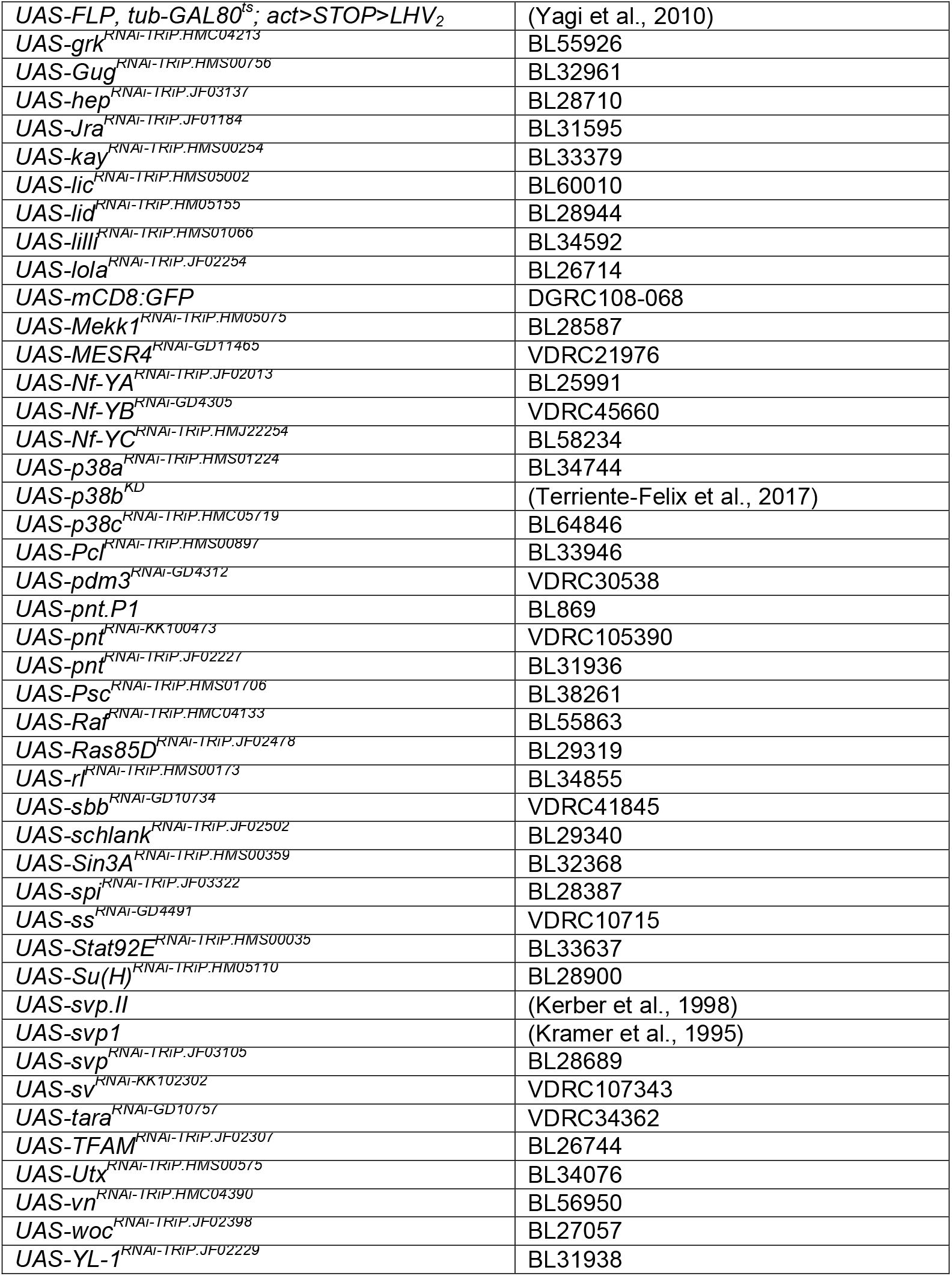
*Drosophila* stocks.

**Table S3.** Primary and secondary antibodies.

| Antibody | Dilution | Source |
| --- | --- | --- |
| Rabbit α-Amos | 1:500 | (zur Lage et al., 2003) |
| Mouse α-Dac | 1:20 | DSHB, Iowa |
| Rat α-Elav | 1:20 | DSHB, Iowa |
| Chicken α-GFP | 1:2000 | Abcam (ab13970) |
| Rabbit α-GFP | 1:1000 | Invitrogen (A6455) |
| Mouse α-nc82 | 1:10 | DSHB, Iowa |
| Rabbit α-Ato | 1:200 | (zur Lage et al., 2003) |
| Rabbit α-Sens | 1:1000 | (Nolo et al., 2000) |
| Mouse α-Svp | 1:10 | DSHB, Iowa |
| Rabbit α-Pon | 1:200 | (Jia et al., 2015) |
| Guinea pig α-Ham | 1:30 | (Moore et al., 2002) |
| Mouse α-Cut | 1:10 | DSHB, Iowa |
| Mouse α-Pros | 1:10 | DSHB, Iowa |
| Rabbit α-Pnt | 1:1000 | (Pascual et al., 2017) |
| Rabbit α-Orco | 1:200 | (Benton et al., 2006) |
| Guinea pig α-Ir8a | 1:300 | (Abuin et al., 2011) |
| Rabbit α-Lush | 1:100 | (Gomez-Diaz et al., 2013) |
| α-DIG-POD | 1:300 | Roche Diagnostics AG (11 207 733 910) |
| α-FITC-POD | 1:300 | Roche Diagnostics AG (1 426 346) |
| α-rabbit Cy3 | 1:1000 | Jackson ImmunoResearch (111-165-144) |
| α-mouse Cy5 | 1:250 | Jackson ImmunoResearch (115-175-166) |
| α-chicken Alexa 488 | 1:1000 | Abcam (ab150169) |
| α-rabbit Alexa 488 | 1:1000 | Invitrogen (A11034) |
| α-rat Alexa 467 | 1:250 | Jackson ImmunoResearch (112-606-062) |
| α-rabbit Cy5 | 1:250 | Jackson ImmunoResearch (111-175-144) |
| α-mouse Cy3 | 1:1000 | Jackson ImmunoResearch (115-165-166) |
| α-guinea pig Cy3 | 1:1000 | Jackson ImmunoResearch (106-166-003) |

**Table S4.**
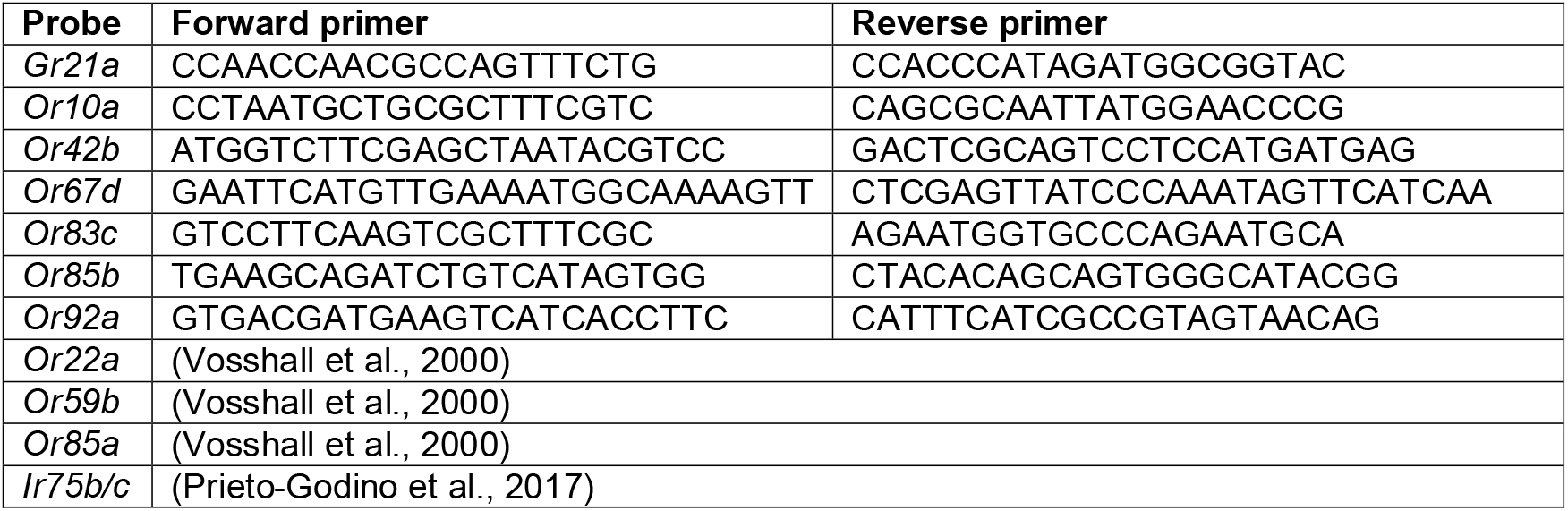
Primers used for generating RNA FISH probes

